# A chronic murine model of pulmonary *Acinetobacter baumannii* infection enabling the investigation of late virulence factors, long-term antibiotic treatments, and polymicrobial infections

**DOI:** 10.1101/2024.09.17.613469

**Authors:** Clay D. Jackson-Litteken, Gisela Di Venanzio, Manon Janet-Maitre, Ítalo A. Castro, Joseph J. Mackel, David A. Rosen, Carolina B. López, Mario F. Feldman

## Abstract

*Acinetobacter baumannii* can cause prolonged infections that disproportionately affect immunocompromised populations. Our understanding of *A. baumannii* respiratory pathogenesis relies on an acute murine infection model with limited clinical relevance that employs an unnaturally high number of bacteria and requires the assessment of bacterial load at 24-36 hours post-infection. Here, we demonstrate that low intranasal inoculums in immunocompromised mice with a *tlr4* mutation leads to reduced inflammation, allowing for persistent infections lasting at least 3 weeks. Using this “chronic infection model,” we determined the adhesin InvL is an imperative virulence factor required during later stages of infection, despite being dispensable in the early phase. We also demonstrate that the chronic model enables the distinction between antibiotics that, although initially reduce bacterial burden, either lead to complete clearance or result in the formation of bacterial persisters. To illustrate how our model can be applied to study polymicrobial infections, we inoculated mice with an active *A. baumannii* infection with *Staphylococcus aureus* or *Klebsiella pneumoniae*. We found that *S. aureus* exacerbates the infection, while *K. pneumoniae* enhances *A. baumannii* clearance. In all, the chronic model overcomes some limitations of the acute pulmonary model, expanding our capabilities to study of *A. baumannii* pathogenesis and lays the groundwork for the development of similar models for other important opportunistic pathogens.

## Introduction

*Acinetobacter baumannii* is a Gram-negative opportunistic pathogen that causes diverse infections including pneumonia, <u>u</u>rinary tract infection (UTI), bone and soft tissue infection, and septicemia (1–5). While becoming an increasingly more common cause of community-acquired infections, *A. baumannii* still primarily causes hospital-acquired infections in critically ill and immunocompromised patients, ∼25% of which are polymicrobial (6–11). These infections are associated with an alarming mortality rate, up to 80% in some populations, largely owing to extremely high rates of multi-drug resistance (8, 12, 13). Notably, *A. baumannii* isolates exhibit the highest rates of multi-drug resistance of all Gram-negative pathogens, leading the World Health Organization to classify the bacterium at its highest priority for research and development of new treatments (13, 14). There is consequently an urgent need to better understand the virulence mechanisms employed by *A. baumannii* to guide the development of novel therapeutic approaches to combat infections.

While *A. baumannii* can cause a variety of infections, it is most commonly associated with pneumonia (4, 15). In fact, *A. baumannii* causes up to 10% of all <u>h</u>ospital-<u>a</u>cquired <u>p</u>neumonia (HAP) cases in the United States, highlighting its importance in clinical settings (16, 17). Despite this, little is known regarding the pathogenesis of this bacterium in the respiratory tract (18). A major hindrance in the ability to investigate *A. baumannii* pneumonia is the lack of available clinically-relevant murine infection models. This is, in large part, due to the low virulence of most strains in immunocompetent mice. This is a shared feature among many pathogens that commonly cause HAP, including *Pseudomonas aeruginosa* and *Staphylococcus aureus*, for which animal models closely mimicking human infection are not available (19, 20). An acute infection model requiring a very high, and rather artificial, inoculum of 10^8^-10^9^ bacteria introduced intranasally or intratracheally is most often used to investigate these pathogens (10, 20, 21). <u>W</u>ild-type (WT) mice will typically either succumb to infection or clear the organism by 72 h, thus requiring early readouts of infection such as bacterial pulmonary titers at 24-36 h. While this model may serve as a useful tool to study pathogenesis early during infection, the quick bacterial clearance does not allow for the study of bacterial virulence mechanisms at later timepoints. Importantly, *A. baumannii* respiratory infection in humans results in an average length of hospital stay of ∼30 days, and this number is much higher in cases caused by multi-drug resistant strains, highlighting the need for a long-term infection model (22, 23). In this pursuit, some laboratories have used antibody or cyclophosphamide treatments to render mice neutropenic (24–30). These treatments initially make mice more susceptible to *A. baumannii* infection, enabling the study of bacterial pathogenesis up to 7 <u>d</u> <u>p</u>ost-infection (dpi) using lower inoculums (∼10^7^ bacteria). However, these models do not achieve stable neutropenia in mice which leads to clearance of infection. To maintain neutropenia over longer periods, multiple injections would be necessary, which can lead to fluctuating neutrophil levels, thereby altering the overall course of disease. A notable caveat to many reports using these currently available immunocompetent and immunocompromised infection models is that older, lab-domesticated strains and non-lung isolates, such as *Ab*19606 and *Ab*17978, are employed, despite the extensive literature demonstrating numerous genotypic and phenotypic differences between these and modern respiratory isolates (31–35). In all, there is an urgent need for alternative infection models to study bacterial pathogenesis during long-term infection by relevant clinical isolates.

Previous reports have used genetically immunocompromised mice to study the role of the host immune response to *A. baumannii* infection. One example is mice carrying a mutation in toll-like receptor 4 (TLR4). TLR4 recognizes the lipid A moiety of bacterial lipo<u>p</u>oly<u>s</u>accharide (LPS) and lipo<u>o</u>ligo<u>s</u>accharide (LOS), the main component of the outer membrane of most Gram-negative bacteria (36–38). The recognition of lipid A by TLR4 triggers a signaling cascade through MyD88- or TRIF-dependent pathways, resulting in increased inflammatory cytokine and type 1 interferon production, respectively (39). The role of TLR4 during *A. baumannii* infection has been examined in murine septicemia, acute pneumonia, UTI, and <u>c</u>atheter-<u>a</u>ssociated <u>UTI</u> (CAUTI) models (40–42). In the acute pneumonia model, Knapp et al. showed that *tlr4* mutant mice had increased *A. baumannii* CFU in the lungs with reduced inflammatory cytokines compared to WT mice (41, 43). Using a bloodstream infection model, Lin et al. demonstrated that WT C3H/FeJ and *tlr4* mutant C3H/HeJ mice had similar bacterial burdens (40, 44). However, all WT mice succumbed to infection by day 4, whereas all *tlr4* mutant mice survived. This could be attributed to WT mice experiencing septic shock associated with increased inflammatory cytokines. Finally, in a UTI model, our laboratory found that *tlr4* mutant C3H/HeJ mice were more susceptible to infection than WT C3H/HeN mice (42). Moreover, we found that C3H/HeJ mice in the UTI model formed small intracellular populations in urothelial cells referred to as *<u>A</u>cinetobacter <u>b</u>aumannii* intracellular reservoirs (ABIRs), which could seed a recurrent infection upon catheterization at higher rates relative to WT mice. In addition to playing a significant role during murine infection, TLR4 is relevant in clinical settings as well. In fact, numerous studies have identified links between *tlr4* polymorphisms and infection outcomes from *A. baumannii* pneumonia (45–47). In all, these studies demonstrate the key role of TLR4 in controlling *A. baumannii* infection and disease progression and highlight the clinical relevance of the associated signaling cascade.

In this work we describe a novel murine model of *A. baumannii* pneumonia that employs *tlr4* mutant mice and low bacterial inoculums (10^5^ bacteria). Using this model, we show that clinically-relevant *A. baumannii* strains can establish chronic infection. We additionally demonstrate that our model enables the discovery of virulence factors not detectable in the acute infection model. Finally, we illustrate how our model can be employed to assess the efficacy of antibiotics over the course of infection and investigate polymicrobial infections.

## Results

### tlr4 mutant mice are susceptible to chronic infection at low inoculums

To assess if *tlr4* mutant mice could serve as permissive hosts for long-term respiratory infection, we performed intranasal inoculations of WT (C3H/HeN) and *tlr4* mutant (C3H/HeJ) mice with high (10^8^) and low (10^5^) inoculums of a modern *A. baumannii* respiratory isolate, G636, and sacrificed groups of mice every 3 days starting at 24 <u>h</u>ours <u>p</u>ost-infection (hpi). At the higher inoculum, WT mice cleared infection by day 4, consistent with previously published results using the acute pulmonary infection model (**Fig. 1A**) (21). *tlr4* mutant mice infected with the higher inoculum also cleared infection relatively early after inoculation, with most mice having no detectable bacteria in the lungs by ∼7 dpi. Strikingly, while WT mice infected with the lower dose of 10^5^ bacteria cleared infection after 1 day, *tlr4* mutant mice maintained detectable bacteria in the lungs out to the latest timepoint tested, 19 dpi (**Fig. 1D**). Despite this long infection course, dissemination to distal organs was rarely detected. This is consistent with the clinical manifestations of non-ventilator *A. baumannii* pneumonia, as less than 20% of patients will develop subsequent bacteremia (48). Notably, by employing confocal microscopy, we were able to visualize bacteria in *tlr4* mutant mice with the low inoculum; at early timepoints (4 hpi and 2 dpi), bacteria were identified inside cells in the <u>b</u>roncho<u>a</u>lveolar lavage fluid (BALF), as well as extracellularly, consistent with our previously published results in the acute infection model (**Fig. S1**) (31).

**Figure 1.**
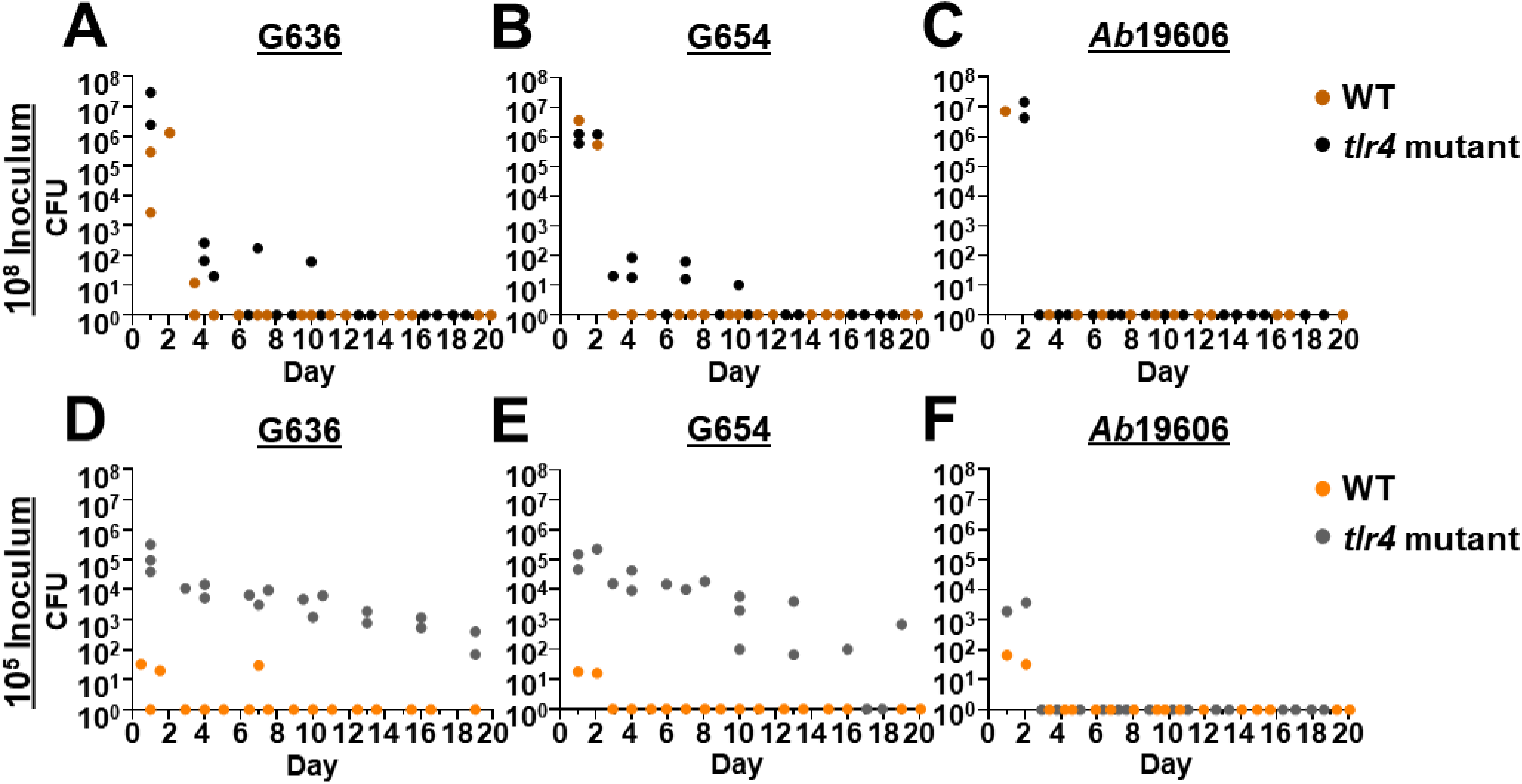
Low inoculums of modern respiratory *A. baumannii* clinical isolates result in chronic lung infection in *tlr4* mutant mice. Groups of C3H/HeN (WT) or C3H/HeJ (*tlr4* mutant) mice were intranasally inoculated with 10^8^ G636 (A), 10^8^ G654 (B), 10^8^ *Ab*19606 (C), 10^5^ G636 (D), 10^5^ G654 (E), or 10^5^ *Ab*19606 (F). Beginning at 24 hpi, groups of mice were sacrificed every three days, and bacteria in the lungs were quantified. Each data point indicates and individual mouse.

We then assessed if a second modern *A. baumannii* respiratory isolate, G654, behaves similarly to G636 (**Fig. 1E**). Again, detectable levels of bacteria were present in the lungs out to 19 dpi with the low inoculum in *tlr4* mutant mice, while WT mice cleared this inoculum within 1 day. At the higher inoculum, G654 was cleared soon after inoculation regardless of TLR4 functionality (**Fig. 1B**). On the contrary, when we tested an older, lab-domesticated urinary isolate that is commonly used to study *A. baumannii* respiratory pathogenesis, *Ab*19606, we found that WT and *tlr4* mutant mice cleared infection after 1 day regardless of inoculum size or mouse background (**Fig. 1C and 1F**). In all, these results indicate that modern *A. baumannii* respiratory isolates can cause infection out to nearly 3 weeks at lower and likely more clinically-relevant inoculums than previously used in the literature. The finding that *Ab*19606 was unable to establish long-term infection further highlights the differences between modern, infection site-specific isolates and lab-domesticated strains (31–35). Importantly, this infection duration with low inoculums of modern respiratory isolates in *tlr4* mutant mice is the longest reported for *A. baumannii* in any animal model to date. We therefore chose to further characterize these conditions as a model to study pulmonary pathogenesis, referred to hereafter as the “chronic respiratory infection model.”

### Lower A. baumannii inoculums result in a decreased immune response in tlr4 mutant mice

Given the unexpected result that *tlr4* mutant mice exhibit chronic infection at lower inoculums, while WT and *tlr4* mutant mice clear infection at higher inoculums, we sought to characterize the host immune response in these different conditions. We intranasally infected groups of WT and *tlr4* mutant mice with 10^5^ or 10^8^ bacteria or mock infected them with <u>p</u>hosphate-<u>b</u>uffered <u>s</u>aline (PBS). Then, at early timepoints of 4 hpi and 2 dpi and a later timepoint of 7 dpi, BALF was collected for immune cell quantification (**Fig. 2**). Regardless of timepoint, inoculum, or mouse background, few significant changes were observed in number of <u>a</u>lveolar <u>m</u>acrophages (AMs) (**Fig. 2A-2C**). In WT mice, the number of <u>p</u>oly<u>m</u>orpho<u>n</u>uclear leukocytes (PMNs) was increased with the higher inoculum relative to lower and mock inoculums at every timepoint (**Fig. 2D-2F**). Additionally, at the higher inoculum, WT mice had increased PMNs relative to *tlr4* mutant mice at every time point in line with previous results (41). Interestingly, at the lower inoculum, while PMN counts trended higher at 4 hpi for WT mice relative to *tlr4* mutant mice, no significant differences were noted between these groups at any timepoint. This result suggests that, despite neutrophil influx being a predominant mechanism of *A. baumannii* clearance, PMN numbers alone may not account for differences in clearance between WT and *tlr4* mutant mice at the lower inoculum (49–55).

**Figure 2.**
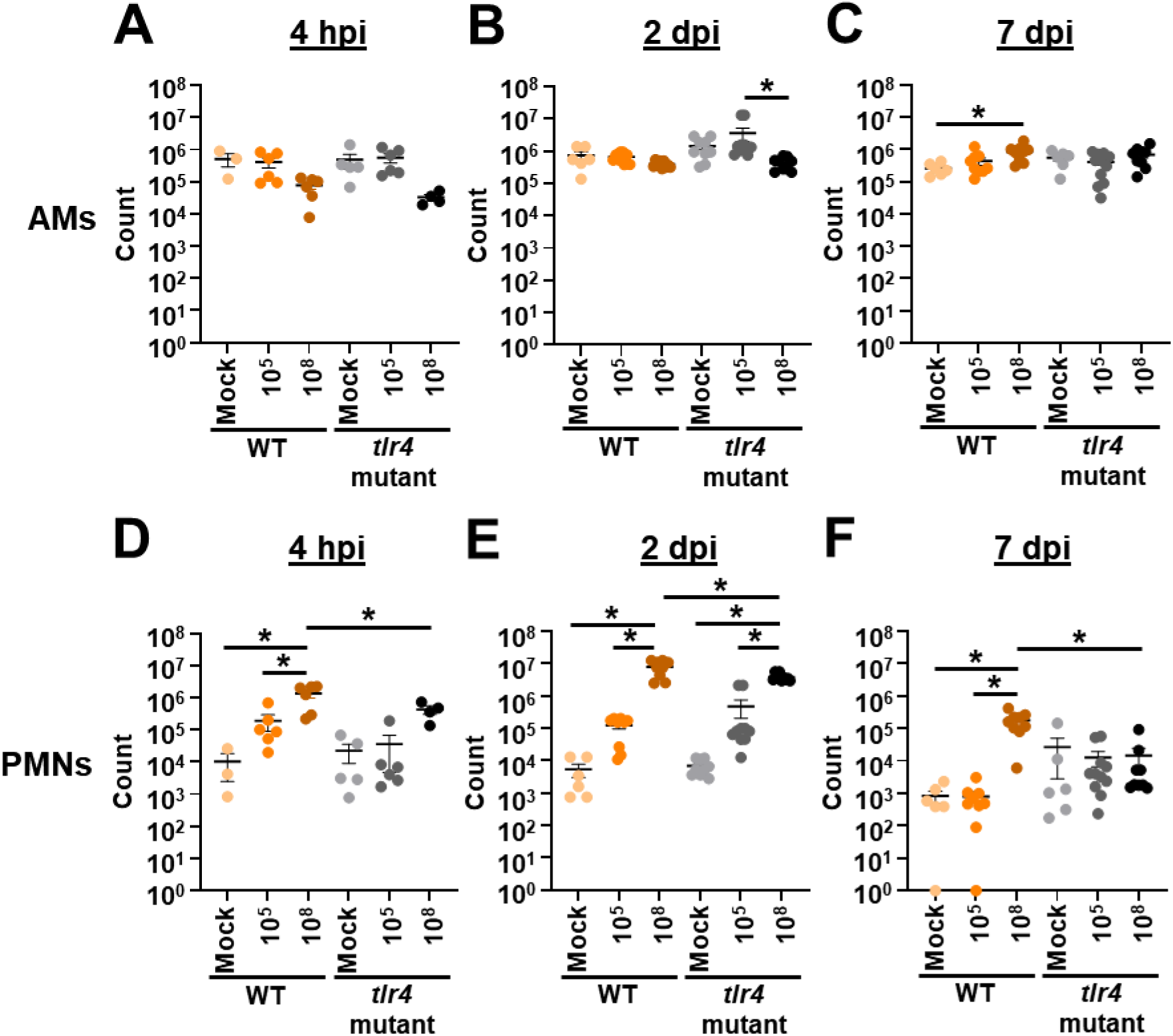
Lower intranasal *A. baumannii* inoculums result in reduced lung neutrophil influx. Groups of C3H/HeN (WT) or C3H/HeJ (*tlr4* mutant) mice were intranasally inoculated with 10^5^ G636, 10^8^ G636, or mock inoculated with PBS. At 4 h (A and D), 2 d (B and E), and 7 d (C and F) pi, <u>a</u>lveolar <u>m</u>acrophages (AMs) (A-C) and <u>p</u>oly<u>m</u>orpho<u>n</u>uclear leukocytes (PMNs) (D-F) in the BALF were enumerated by flow cytometry. Shown are pooled results from at least two independent experiments, and each data point represents an individual mouse. The horizontal line represents the mean, and the <u>s</u>tandard <u>e</u>rror of the <u>m</u>ean (SEM) is indicated by error bars. \**P* < 0.05; two-way <u>an</u>alysis <u>o</u>f <u>va</u>riance (ANOVA), Tukey’s test for multiple comparisons.

To further evaluate the host response, we quantified 13 common inflammatory cytokines in the BALF (**Table S1**). At the higher inoculum, WT and *tlr4* mutant mice exhibited significantly increased levels of IL-1α, IFN-γ, TNF-α, MCP-1, IL-1β, IL-6, and IL-17A early during infection relative to the lower inoculum while levels dissipated by 7 dpi, consistent with bacterial clearance (See **Fig. 1A**). WT mice infected with the high inoculum had significantly increased levels of IFN-β relative to *tlr4* mutant mice at 4 hpi and increased levels of IL-1α, IFN-γ, TNF-α, MCP-1, IL12-p70, IL-1β, IL-6, IL-27, and IL17A at 2 dpi, likely leading to the earlier clearance observed. At the lower inoculum, although WT mice clear infection within nearly 24 h and *tlr4* mutant mice maintain infection out to at least three weeks, minimal significant differences in inflammatory cytokines were observed (See **Fig. 1D**). In fact, the only significant difference noted was the increased levels of GM-CSF at 4 hpi in WT mice relative to *tlr4* mutant mice. Other inflammatory cytokines that trended higher at early timepoints in WT mice at the lower inoculum include the inflammasome-associated cytokines IL-1α and IL-1β, as well as TNF-α and IL-6. These elevated levels of inflammatory cytokines early during infection in WT mice could possibly account for the earlier clearance. Later during infection, however, *tlr4* mutant mice had elevated, albeit not significantly higher, amounts of TNF-α, IL-1α, and IL-6 relative to WT mice, consistent with persistent infection.

### The chronic respiratory infection model results in lung pathology

We next assessed if the chronic respiratory infection model is associated with lung pathology. *tlr4* mutant mice were infected with 10^5^ G636 or mock infected with PBS, and mice were sacrificed at 4 hpi, 2 dpi, 7 dpi, 14 dpi, and 21 dpi. Lungs were then sectioned, stained with <u>h</u>ematoxylin and <u>e</u>osin (H&E), and scored for pathological changes as previously described (**Fig. 3**, **Fig. S2**, and **Fig. S3**) (56). The chronic model resulted in significant increases in alveolitis, peribronchiolitis, smooth muscle hypertrophy, squamous epithelium metaplasia, and formation of <u>b</u>ronchus-<u>a</u>ssociated lymphoid tissue (BALT) relative to mock-treated mice (**Fig. 3** and **Fig S2**). Significant changes in goblet cell hyperplasia and fibrosis were not detected (**Fig. S3**). Of note, significant signs of disease were detected out to 14 dpi, and, even at 21 dpi, infected mice showed trends toward increased lung damage relative to mock-infected mice. In all, histopathological analyses revealed that the chronic respiratory infection model results in sustained lung damage, indicative of chronic infection.

**Figure 3.**
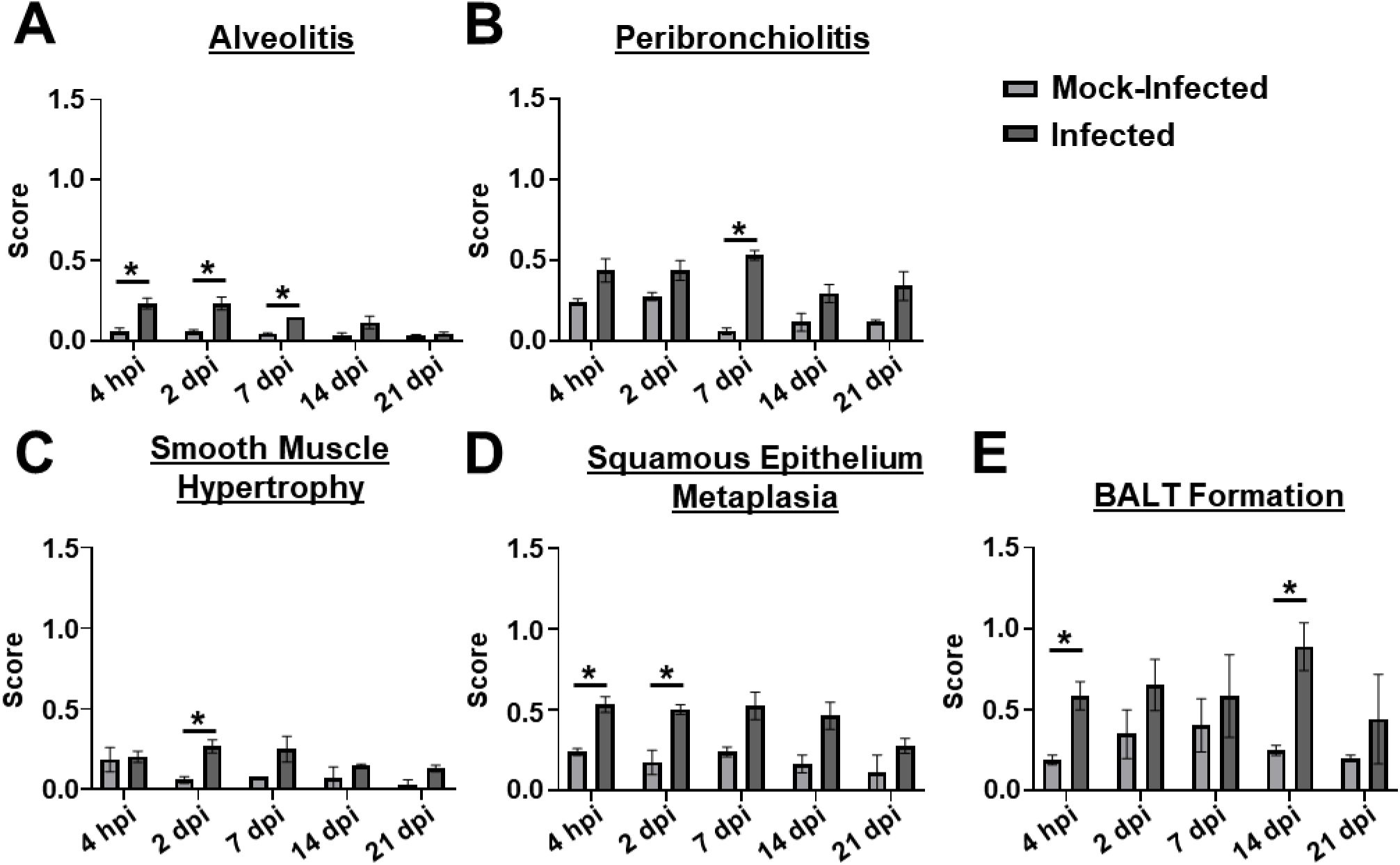
The chronic respiratory infection model results in lung pathology. Groups of C3H/HeJ (*tlr4* mutant) mice were inoculated with 10^5^ G636 or mock-inoculated with PBS, and at 4 hpi, 2 dpi, 7 dpi, 14 dpi, and 21 dpi, lungs slices were prepared, H&E stained, and scored for alveolitis (A), peribronchiolitis (B), smooth muscle hypertrophy (C), squamous epithelium metaplasia (D), and BALT formation (E). The mean is shown on the graph, and the SEM is indicated by error bars. \**P* < 0.05, Unpaired Student’s *t*-test.

### InvL is a critical virulence factor for long-term infection

The acute pulmonary infection model has been widely used to characterize *A. baumannii* virulence factors (21). While the acute model is valuable for identifying bacterial proteins required at early timepoints, these mice clear infection within 3-4 dpi, not allowing for the identification of factors required for prolonged infection. As a proof of principle, we sought to determine if the chronic respiratory infection model could identify proteins required for bacterial persistence in the lungs. We hypothesized that prolonged adherence to respiratory epithelium would be required for persistence, so we first tested individual mutants lacking previously identified *A. baumannii* adhesins (Bap, Ata, FhaBC, and InvL) for attenuation in the chronic infection model (**Fig. S4**) (57–66). *tlr4* mutant mice were infected with 10^5^ G636 WT or mutant bacteria, and mice were sacrificed at 1 or 14 dpi for lung <u>c</u>olony-forming <u>u</u>nits (CFU) quantification. This experiment indicated a possible role for InvL in long-term infection as mice began to clear bacteria in the lungs by 14 dpi. To further confirm the importance of InvL for bacterial persistence, we performed more extensive analyses with G636 WT, *invL* mutant (Δ*invL*), and complemented *invL* mutant (*invL*^+^) strains in the chronic infection model, sacrificing mice at 1, 7, 14, and 21 dpi to quantify CFU in the lungs (**Fig. 4A-D**). Early during infection, the Δ*invL* mutant exhibited only a modest defect. However, at later timepoints the infection defect became more pronounced, as some mice cleared the bacteria as early as 7 dpi. By 21 dpi, all but two mice had cleared the Δ*invL* mutant, while the majority of mice infected with the WT strain still had detectable bacteria in their lungs. Genetic complementation partially rescued this defect at these later timepoints, as no significant difference was detected between WT and complemented strains.

**Figure 4.**
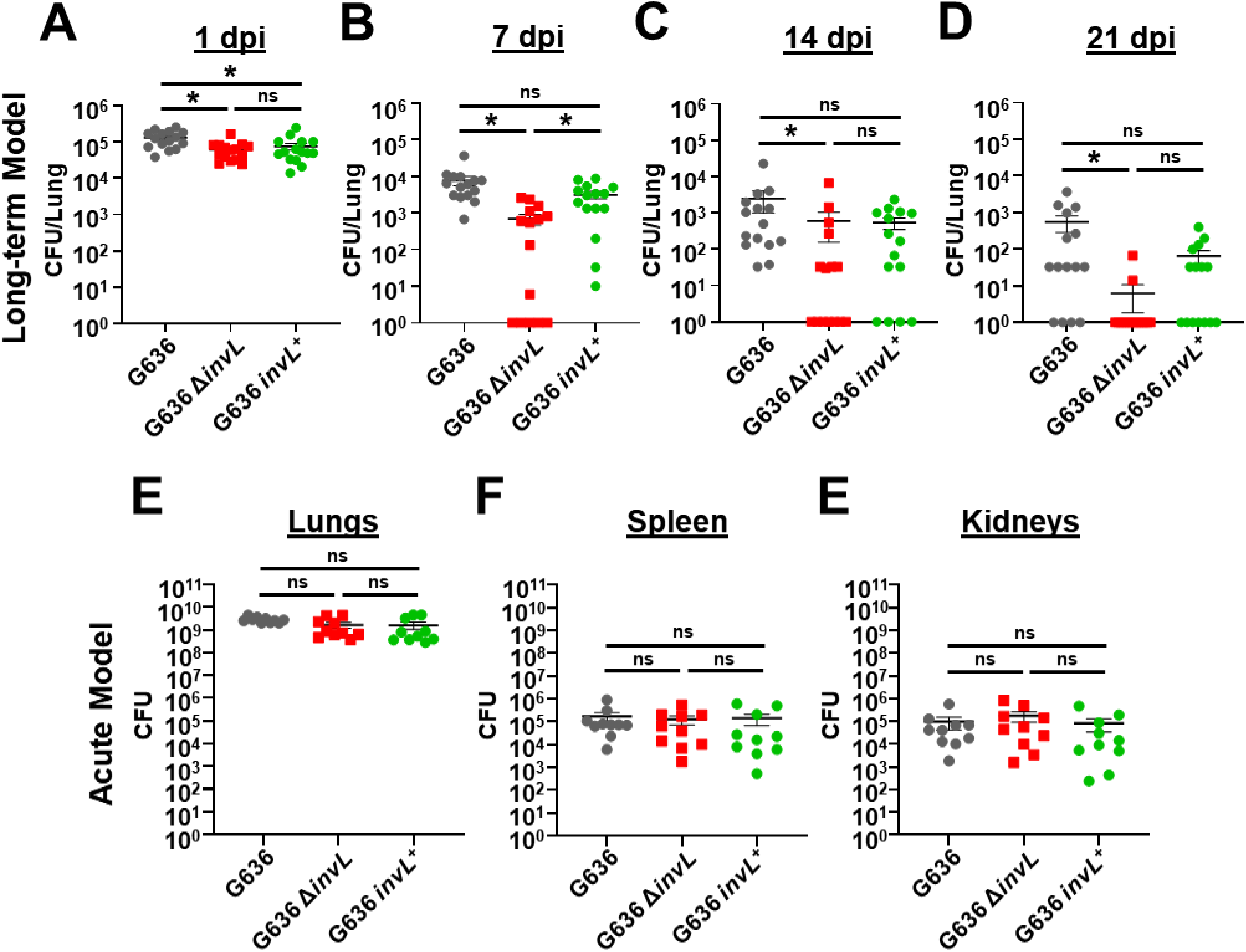
InvL is a critical virulence factor for long-term respiratory infection, but dispensable in the acute infection model. C3H/HeJ (*tlr4* mutant) mice were infected with 10^5^ G636, G636 Δ*invL*, or G636 *invL*^+^. Groups of mice were then sacrificed at 1 dpi (A), 7 dpi (B), 14 dpi (C), and 21 dpi (D), and CFU in the lungs were quantified. Shown are the results from 3 independent experiments. For the acute infection model, groups of C57BL/6 mice were infected with 10^9^ G636, G636 Δ*invL*, or G636 *invL^+^*. 24 hpi, mice were sacrificed, and CFU in the lungs (A), spleen (B), and kidneys (C) were enumerated. Each data point represents an individual mouse, the horizontal line represents the mean, and the SEM is indicated by error bars. Shown are the results of 2-3 independent experiments. \**P* < 0.05; Kruskal-Wallis *H* test with Dunn’s test for multiple comparisons; ns = not significant.

We next compared results from the chronic respiratory infection model to the acute infection model. We infected C57BL/6 mice with 10^9^ G636 WT, Δ*invL*, or *invL*^+^ strains. 24 hpi, mice were sacrificed, and CFU in the lungs, spleens, and kidneys were quantified (**Fig. 4E-F**). As opposed to results seen in the chronic infection model, the Δ*invL* mutant had no significant defect in bacterial load in the lungs. Additionally, no defect was noted in dissemination to the spleen and kidneys, indicating that InvL is dispensable in the acute infection model. In all, these results highlight the differences in required bacterial genes between these disparate pulmonary infection models and show the importance of continuing to explore models that can better approximate clinical disease. Additionally, these experiments establish InvL as the first known *A. baumannii* virulence factor required for long-term infection.

### The chronic infection model can be used to study the outcome of antibiotic treatment

The acute pulmonary infection model has been employed extensively to assess effects of antibiotic treatment (21). However, this model only allows us to estimate the efficacy of antibiotics by measuring the initial reduction in the bacterial burden at 24-36 hpi due to rapid bacterial clearance by the host. A clear limitation of this model is that it does not inform if bacterial infection is cleared, or if persistent bacteria remain in the lung. The chronic respiratory infection model therefore represents a novel platform that could be used to track the kinetics of *A. baumannii* clearance due to antibiotic treatment. As a proof of principle, we assessed the effect of tigecycline, colistin, and imipenem in the chronic model with strains G636 and G654 at antibiotic concentrations similar to those previously used to determine treatment efficacy in mice (**Fig. 5A-B** and **Fig. S5A-D**) (67–73). We additionally assessed the effect of apramycin, a drug with demonstrated efficacy and safety in mice that is currently in Phase I clinical trials for use in humans (**Fig. 5C-D**) (74–76). <u>M</u>inimum inhibitory <u>c</u>oncentrations (MICs) for G636 and G654 for these and other commonly used antibiotics are listed in **Table S2**. Colistin was ineffective for both strains in the acute infection model, while initial reductions in CFU were noted in the chronic infection model (**Fig. S5A-B**). However, bacterial numbers appeared to stabilize over time in the chronic infection model, consistent with the development of bacterial persisters (discussed below). Imipenem showed limited efficacy against both strains in both models (**Fig. S5C-D**), as expected given the strains’ resistance *in vitro* (**Table S2**). At 24 hpi, tigecycline and apramycin treatment resulted in initial reductions in CFUs in both the chronic and the acute infection models relative to PBS-treated mice (**Fig. 5A-D**). However, the chronic model enabled us to differentiate the efficacy of both antibiotics at later times. Apramycin treatment ultimately led to clearance after 3-5 days, demonstrating the efficacy of this antibiotic. However, with tigecycline treatment, although there were initial reductions in CFU, bacterial numbers leveled out over time indicative of treatment failure. The behavior of bacteria in presence of tigecycline over time is consistent with the development of persisters. Notably, the efficacy of tigecycline and apramycin against *A. baumannii* cannot be distinguished at 24 hpi. These results indicate that the chronic model can be used to determine outcome of infection with therapeutic intervention, a significant advantage over the currently employed acute infection model.

**Figure 5.**
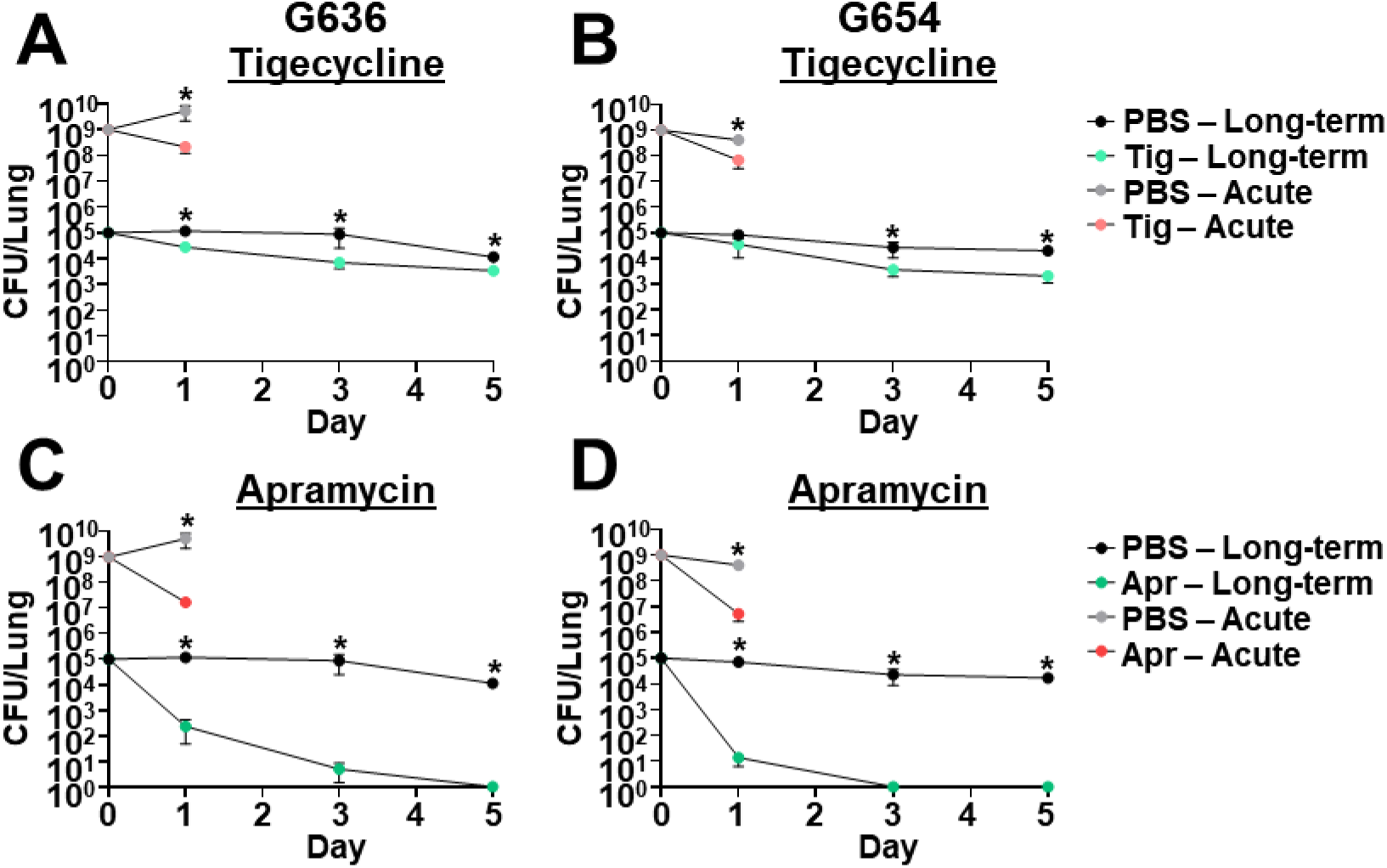
The chronic respiratory infection model can be used to study outcomes of antibiotic treatment. Groups of C3H/HeJ (*tlr4* mutant) mice were infected with 10^5^ G636 (A, C) or 10^5^ G654 (B, D) and sacrificed at 1, 3, and 5 dpi (long-term). Additionally, groups of C57Bl/6 mice were infected with 10^9^ G636 (A, C) or 10^9^ G654 (B, D) and sacrificed at 24 hpi (acute). Mice in both infection models were treated intraperitoneally treated with PBS or 100 mg/kg <u>tig</u>ecycline (tig) every 12 h (A, B) or PBS or 500 mg/kg <u>apr</u>amycin (apr) every 12 h (C, D) with all treatments beginning 4 hpi. At each timepoint, CFU were quantified in the lungs. Shown are the results from at least two independent experiments, each data point represents an individual mouse, the horizontal line represents the mean, and the SEM is represented by error bars. \**P* < 0.05; Mann-Whitney *U* test.

### Use of the chronic infection model to study bacterial co-infections reveals that Staphylococcus aureus exacerbates ongoing A. baumannii infection while Klebsiella pneumoniae leads to earlier clearance

Approximately 25% of *A. baumannii* pulmonary infections are polymicrobial, and two of the most commonly co-infecting pathogens are *Staphylococcus aureus* and *Klebsiella pneumoniae* (9). We thus sought to assess the impact of secondary infections with these two bacteria on the outcome of *A. baumannii* infection in the context of the chronic respiratory infection model. For these experiments, we first established a primary *A. baumannii* infection by inoculating *tlr4* mutant mice with 10^5^ CFU of strain G636. Following 14 days of *A. baumannii* infection, we inoculated mice with 5 x 10^7^ CFU of *S. aureus* strain Newman or *K. pneumoniae* strain TOP52, mock-treated mice with PBS, or left mice untreated. One and two days post-secondary infection, mice were sacrificed, and bacterial CFU were quantified in the lungs, spleens, and kidneys (**Fig. 6**, **Fig. S6**, and **Fig. S7**). Secondary infection with *S. aureus* led to a resurgence of *A. baumannii* CFU in the lungs of many mice, though the overall mean CFU in these mice were not significantly different from mock-infected and untreated groups (**Fig. 6A** and **Fig. 6D-E**). *A. baumannii* were also identified in the spleens and kidneys of some mice that received the secondary *S. aureus* infection, even though *A. baumannii* bacteremia rarely occurs in the context of this chronic respiratory infection model (**Fig. S6**). Additionally, *S. aureus* trended toward increased numbers in the lungs, spleens, and kidneys in the context of polymicrobial infection with *A. baumannii* relative to monomicrobial infection (**Fig. S7A-C**). Notably, two mice succumbed to *A. baumannii*-*S. aureus* polymicrobial infection ∼24 hpi following the secondary inoculation, an outcome that did not occur with monomicrobial infection with either bacterium. Contrarily, secondary infection with *K. pneumoniae* significantly decreased *A. baumannii* CFU in the lungs relative to mock-infected and untreated groups (**Fig. 6A** and **Fig. 6D-E**). Additionally, polymicrobial infection with *A. baumannii* and *K. pneumoniae* resulted in significantly reduced *K. pneumoniae* CFU recovered in the lungs, spleens, and kidneys of mice relative to *K. pneumoniae* monomicrobial infection (**Fig. S7D-E**). Although understanding the interactions between these bacteria is beyond the scope of this work, these experiments indicate that *S. aureus* exacerbates *A. baumannii* infection, while *K. pneumoniae* attenuates infection in the context of the chronic infection model. Additionally, these results demonstrate the ability of the model to be used to study longer-term aspects of polymicrobial interactions that were not previously able to be done with the acute infection model.

**Figure 6.**
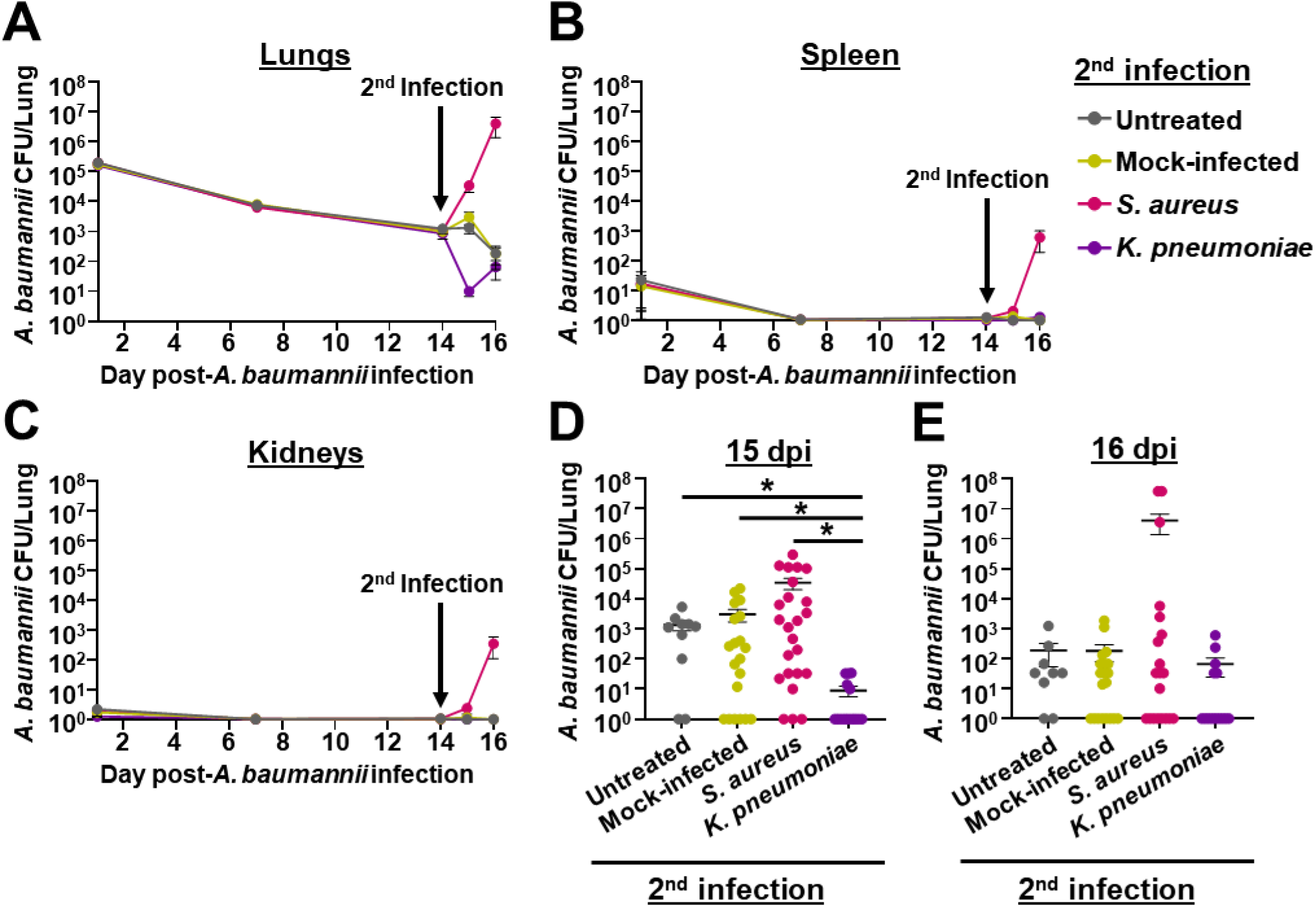
Bacterial secondary infection alters the course of chronic *A. baumannii* pneumonia. C3H/HeJ (*tlr4* mutant) mice were intranasally inoculated with 10^5^ G636, and groups of mice were sacrificed at 1, 7, and 14 dpi. At 14 days post-*A. baumannii* infection, groups of mice were either not inoculated (untreated), inoculated with PBS (mock-infected), infected with *S. aureus*, or infected on *K. pneumoniae*. Subsequently, on days 15 and 16 post-*A. baumannii* infection (1 and 2 days post-secondary infection), mice were sacrificed, and *A. baumannii* CFU were quantified in the lungs (A, D, E), spleen (B), and kidneys (C). In panels A, B, and C, each data point represents the mean, the SEM is represented by error bars, and the limit of detection is indicated by the dashed line. In panels D and E, each data point represents an individual mouse, the horizontal line represents the mean, and the SEM is indicated by error bars. Shown are results from at least 2 independent experiments. \**P* < 0.05; Kruskal-Wallis *H* test with Dunn’s test for multiple comparisons.

## Discussion

*A. baumannii* has emerged as a significant cause of nosocomial pneumonia and is of major clinical importance due to its extremely high rates of multidrug resistance (8, 12, 13). Despite this, our understanding of *A. baumannii* respiratory pathogenesis is hindered by a shortage of clinically relevant infection models. Here, we aimed to address this significant gap in the field by developing a novel respiratory infection model. In this pursuit, we found that, at likely more clinically-relevant inoculums, *tlr4* mutant mice maintain long-term respiratory infections by *A. baumannii*. We then demonstrate the versatility of this model which enabled i) the identification of a bacterial virulence factor required for long-term respiratory infection, which is not required in acute models, ii) the study of kinetics of bacterial clearance upon treatment with clinically-relevant antibiotics, and iii) the exploration of the impact of secondary infections with two commonly co-isolated respiratory pathogens.

In this study, we found that InvL is required for chronic infection, and, more importantly, at the later stages of infection. However, InvL was dispensable in the context of the acute infection model. There are multiple possible reasons for this discrepancy. First, the massive bacterial dose required for the acute infection model may mask potential defects that can now be detected with a smaller, more clinically-relevant inoculum. This is unlikely, as WT and *invL* mutant bacteria behave similarly at early time points in our model. An alternative reason could be that adhesins required early during infection/interaction with the healthy airway differ from those required during persistent interaction with a more inflamed or damaged airway. It is well-established that the airway <u>e</u>xtra<u>c</u>ellular <u>m</u>atrix (ECM) is altered by bacterial infection, lung damage, and/or inflammation (91, 92). Long-term lung damage and inflammation results in increased fibronectin, collagen, laminin, and fibrinogen in the ECM (93–98). Moreover, specific pathogens elicit different inflammatory responses, resulting in distinct changes to the lung ECM. For example, in an acute mouse model of pneumonia, *Pseudomonas aeruginosa* induces versican deposition in the lungs, while *Escherichia coli* induces robust versican and hyaluronan deposition (99, 100). We previously showed that InvL can bind α5β1 integrin, collagen V, and fibrinogen (66). However, whether *A. baumannii* infection or the associated inflammation induces production of these protein(s) during pulmonary infection is unknown. Future work will investigate this possibility, as well as assess which InvL-host protein interactions are essential for chronic infection.

Herein, we demonstrate the potential to use the chronic respiratory infection model to study the efficacy of antibiotic treatments over time. One intriguing finding from these experiments is that with antibiotics such as colistin and tigecycline, an initial decrease in CFU (∼10-100 fold) recovered from the lungs at 1 dpi was observed. However, following this decrease, the number of bacteria in the lungs appeared to stabilize over time. It is tempting to speculate that this is the result of the formation of bacterial persisters, defined as bacterial cells that become tolerant to antibiotics despite undergoing no genetic changes (101–103). Importantly, the commonly used acute infection model does not allow for the study of bacterial persisters due to the short time course of the model. Given that persister cells represent a major cause of treatment failure and chronic infection, the chronic infection model presented here represents a unique platform that is desperately needed to understand this aspect of *A. baumannii* pathogenesis. Furthermore, our model offers new possibilities to study efficacy of novel antibiotics in murine models before committing to expensive clinical trials.

While a significant portion of *A. baumannii* infections are polymicrobial, the acute infection model has limitations for use with polymicrobial infections. First the quick clearance of the bacteria usually only allows inoculation at a single timepoint, thus not enabling investigation of secondary infections. Second, the high required infectious dose often means that typical inoculums for bacteria used in these experiments must be adjusted, so mice do not succumb to infections at early timepoints. Here, we applied the chronic respiratory infection model to assess the result of secondary infection with two pathogens commonly co-isolated with *A. baumannii*, *S. aureus* and *K. pneumoniae*. We found opposite results with these different bacteria; *S. aureus* secondary infection trended toward exacerbation of *A. baumannii* infection, while *K. pneumoniae* secondary infection led to reduced *A. baumannii* numbers. The potential synergism of *A. baumannii* and *S. aureus* in the chronic infection model aligns with previous reports. For example, using a Tn-Seq-based approach, Li et al. demonstrated that the 49% of genes required by *S. aureus* for monomicrobial infection in a murine systemic infection model became non-essential upon *A. baumannii* co-infection (104). Another recent report showed that *S. aureus* can support *A. baumannii* growth *in vitro* by providing acetoin as a carbon source (105).

Although we found that *K. pneumoniae* secondary infection led to reduced *A. baumannii* numbers in the lungs in the chronic infection model, one study has shown that *K. pneumoniae* could cross-feed *A. baumannii* through products of sugar fermentation *in vitro* and demonstrated that co-infection led to reduced survival of *Galleria mellonella* relative to monomicrobial infection with either pathogen (106). This, in part, shows that these two bacteria can have beneficial interactions. There are two potential reasons however for the reduction of CFU for both bacteria in the context of the chronic infection model reported here; i) bacterial competition or ii) the host response to the secondary infection. Regarding bacterial competition, there have been several lines of evidence pointing to direct bacterial killing between diverse *A. baumannii* and *K. pneumoniae* strains mediated by the type VI secretion system (107–110). In addition to direct killing, this bacterial competition could be indirect as well, as both *A. baumannii* and *K. pneumoniae* may be competing for similar nutrients in the lung microenvironment. With respect to the immune response, a difference between this work and the above study is that the microenvironment encountered in the mammalian lung is not perfectly modeled by the wax moth (111). Our results may therefore be the result of TLR4-independent host response elicited by the combination of both bacteria that is not recapitulated by a *G. mellonella* model. While understanding the precise mechanism behind the *in vivo* interactions between *A. baumannii* and commonly co-isolated pathogens is outside the scope of the current study, these results highlight the practicality of applying the chronic respiratory infection model to better understand polymicrobial infections.

In this study, we have validated several different uses for the chronic respiratory infection model. However, there are also other potential uses for this model that were not previously investigable. For example, we can now perform experiments differentiating between virulence factors required for establishment of infection and factors required for maintenance of infection, assessing bacterial evolution during long-term infection, investigating changes in the pulmonary microbiome due to infection over time, and analyzing the long-term outcomes of novel therapies such as newly developed phage cocktails. Additionally, while this model was initially developed to study *Acinetobacter* respiratory infections, it has the potential to be applied to research with other respiratory pathogens in cases where suitable animal models are lacking. In all, this work describes the longest-term infection model available to investigate *A. baumannii* host-pathogen interactions to date, which will ultimately aid in the development of novel therapeutics to combat infection by this increasingly multidrug-resistant bacterium.

## Materials and Methods

### Bacterial plasmids, strains, and growth conditions

Plasmids and strains used in this study are detailed in **Table S3**. Bacterial cultures were grown at 37°C in Lennox broth/agar supplemented with 10 μg/mL chloramphenicol, 50 μg/mL apramycin, 100 μg/mL ampicillin, 50 μg/mL kanamycin, 10 μg/mL tetracycline, or 10% sucrose when appropriate.

### Murine pneumonia models

All animal experiments were approved by the Washington University Animal Care and Use Committee, and we have complied with all relevant ethical regulations. The acute pneumonia model was performed similar to previously described experiments (21, 112). Briefly, overnight cultures were subcultured at a 1:200 dilution and grown shaking at 37°C for 3 h to mid-exponential growth phase. Six- to eight-week-old female C57BL/6 mice (Charles River Laboratories, Wilmington, MA) anesthetized with 4% isoflurane were intranasally inoculated with 10^9^ CFU that were twice-washed in PBS. At 24 hpi, mice were sacrificed, and CFU in the lungs, spleen, and kidneys were quantified by serial dilution plating the homogenized organs. For experiments with C3H/HeN (Envigo International Holdings, Indianapolis, IN) and C3H/HeJ (Jackson Laboratory, Bar Harbor, ME) mice, *A. baumannii*, *S. aureus*, and *K. pneumoniae* inoculums were prepared and mice were intranasally inoculated as described above, with the exception that inoculums of 10^5^ and 10^8^ CFU were used for *A. baumannii*, and 5 x 10^7^ CFU was used for *S. aureus* and *K. pneumoniae*. Following, at the indicated timepoints, mice were sacrificed and bacteria in the lungs, spleen, and kidneys were quantified as described above. For co-infections, *A. baumannii* was distinguished from *S. aureus* and *K. pneumoniae* by plating on LB agar supplemented with 10 μg/mL chloramphenicol. For antibiotic treatment experiments the indicated mice were treated intraperitoneally with PBS or 100 mg/kg tigecycline every 12 h, PBS or 5 mg/kg colistin every 8 h, PBS or 500 mg/kg apramycin every 12 h, or PBS or 100 mg/kg imipenem every 12 h with all treatments beginning 4 hpi. Antibiotics for intraperitoneal treatments were dissolved in PBS, and the injection volume was 100 μl.

### Flow cytometry

Flow cytometry was performed similarly to previously described methods (31). Briefly, BALF samples were collected in PBS supplemented with 1 mM EDTA, and cells were collected by centrifugation at 300 x *g* for 5 min. Cells were then resuspended in Pharm Lyse Buffer (BD Biosciences, Franklin Lakes, NJ) and incubated for 3 min at room temperature to lyse red blood cells. Cells were subsequently washed in fluorescence-<u>a</u>ctivated <u>c</u>ell <u>s</u>orting (FACS) buffer (PBS supplemented with 1% heat inactivated fetal bovine serum and 0.1% sodium azide) and blocked with TruStain FcX PLUS (BioLegend, San Diego, CA) for 15 min at 4°C. Samples were then stained with anti-CD45-BV605 (BioLegend), anti-CD11c-APC (BioLegend), anti-SiglecF-PerCP5.5 (BioLegend), and anti-Ly6G-BV421 (Biolegend) for 30 min at 4°C. Following, cells were washed in FACS buffer and fixed in 2% <u>p</u>araform<u>a</u>ldehyde (PFA). Samples were read on a LSR II Fortessa cytometer (BD Biosciences) or an Aurora cytometer (Cytek Biosciences, Fremont, CA). Total cell counts in the BALF were calculated using Precision Count Beads (BioLegend) according to the manufacturer’s instructions.

### Antibiotic protection assays

Antibiotic protection assays were performed as previously described (31). To determine the number of total and intracellular bacteria present in BALF from *A*. *baumannii* infected mice, two 500 μl aliquots of lavage fluid were centrifuged at 4100 x *g* for 5 min. Pelleted cells were resuspended in warm <u>D</u>ulbecco’s <u>M</u>odified <u>E</u>agle <u>M</u>edium (DMEM) (total bacteria) or DMEM with colistin (50 μg/mL) (intracellular bacteria) and incubated for 1h at 37°C. Samples were then washed three times with PBS and lysed with 500 μL of Triton X-100 (0.05%). CFUs were determined by serial dilutions of the bacterial suspensions. The remaining lungs following BALF collection were also homogenized, and CFUs were quantified by serial dilution plating.

### Cytospin of BALF cells

Cytospin of BALF cells was performed similar to previously described work (31). BALF samples were centrifugated at 300 x *g* for 5 min, and the pellets were resuspended in 1 mL Pharm Lyse Buffer (BD Biosciences) and incubated for 5 min on ice to lyse red blood cells. 9 mL of PBS was added to stop the lysis, viability was determined using Trypan Blue solution (Sigma-Aldrich, St. Louis, MO), and cells were counted using the TC20 Automated Cell Counter (Bio-Rad Laboratories, Hercules, CA). Samples were centrifugated at 300 x *g* for 6 min onto CytoPro Poly-L-Lysine Coated Microscope Slides (ELITechGroup Inc., Logan, UT) using a Cytospin Cytocentrifugue (Fisher Scientific, Hampton, NH). The slides were air-dried overnight at 4°C and fixed in 4% PFA for 30 min at room temperature. Samples were incubated with permeabilizing and blocking solution (PBS supplemented with 0.1% saponin, 0.5% bovine serum albumin, and 10% heat inactivated fetal bovine serum). Cells were stained with Alexa Fluor 555 Phalloidin (Cell Signaling Technology, Danvers, MA) and 4’,6-<u>Dia</u>midino-2-<u>p</u>henylindole dihydrochloride (DAPI) solution (Invitrogen) for 1 h at 37°C. After staining, the samples were rinsed with washing solution [PBS supplemented with 0.1% saponin and 0.5% <u>b</u>ovine <u>s</u>erum <u>a</u>lbumin (BSA)], and then rinsed with water and mounted on a coverslip in ProLong Gold Antifade Mountant (Invitrogen).

### Confocal microscopy

Confocal microscopy was performed as previously described (31, 66). Microscopy slides were analyzed with a Zeiss LSM880 laser scanning confocal microscope (Carl Zeiss AG, Oberkochen, Germany) equipped with 405nm diode, 488nm Argon, 543nm HeNe, and 633nm HeNe lasers. A Plan-Apochromat 63X DIC objective and ZEN black 2.1 SP3 software were used for image acquisition. Images were analyzed using ImageJ software (National Institutes of Health, Bethesda, MD) (113).

### Cytokine analysis

BALF was collected and centrifuged at 300 x *g* for 5 min. Supernatant containing cytokines was then collected and frozen at −20°C until the analysis was performed. Cytokine levels were determined using the LEGENDplex Mouse Inflammation Panel (13-plex) with V-bottom Plate (BioLegend) according to the manufacturer’s instructions. Samples were read using an Aurora cytometer (Cytek Biosciences).

### Histopathology of lung slices

Lung slices were prepared, stained, and scored as previously described by Castro et al. (71). Briefly, lungs were perfused with PBS, inflated with <u>o</u>ptimal <u>c</u>utting temperature (OCT) compound (Fisher Scientific) diluted in 4% PFA at a 1:1 ratio, snap-frozen, and stored at −80°C until sectioning. For histology imaging, 4 μm tissue sections were stained with H&E and imaged with a ZEISS Axioscan 7 Microscope Slide Scanner (Carl Zeiss AG). Lung tissues were blindly scored on a scale of 0 to 3 for alveolitis, peribronchiolitis, smooth muscle hypertrophy, squamous epithelium metaplasia, BALT formation, goblet cell hyperplasia, and fibrosis. Area affected was quantified, multiplied by previously defined intensity scores, and the resulting weighted scores are reported.

### Generation of constructs and strains used in this study

Primers used in this study are listed in **Table S4**. DNA fragments were assembled using either the In-Fusion HD EcoDry Cloning Kit (TaKaRa Bio, Mountain View, CA) or NEBuilder HiFi DNA Assembly Master Mix (New England Biolabs, Ipswich, MA). To generate the vector for generation of the *invL* mutational construct, pEX18Tc was amplified without the tetracycline resistance cassette (primers: 5’ pEX18 marker swap and 3’ pEX18 marker swap), the apramycin resistance cassette was amplified from pKD4-Apr (primers: 5’ Apr for pEX18Ap and 3’ Apr for pEX18Ap), and the amplicons were assembled, generating pEX18Ap (114, 115). The pEX18Ap mutational constructs were then made by amplifying the pEX18Ap vector (primers: 5’ pEX18Tc and 3’ pEX18Tc), a ∼1000 bp region upstream of the genes of interest (*invL*KO primers: 5’ F1 G636 *invL*KO and 3’ F1 G636 *invL*KO; *bap*KO primers: 5’ F1 G636 *bap*KO and 3’ F1 G636 *bap*KO; *ata*KO primers: 5’ F1 G636 *ata*KO and 3’ F1 G636 *ata*KO; *fhaBC*KO primers: 5’ F1 G636 *fhaBC*KO and 3’ F1 G636 *fhaBC*KO), and a ∼1000 bp region downstream of the genes of interest (*invL*KO primers: 5’ F2 G636 *invL*KO and 3’ F2 G636 *invL*KO; *bap*KO primers: 5’ F2 G636 *bap*KO and 3’ F2 G636 *bap*KO; *ata*KO primers: 5’ F2 G636 *ata*KO and 3’ F2 G636 *ata*KO; *fhaBC*KO primers: 5’ F2 G636 *fhaBC*KO and 3’ F2 G636 *fhaBC*KO), followed by assembly of these amplicons. Mutational constructs were then transformed into G636, and strains with the integrated plasmid were selected for by apramycin treatment. Counterselection for double crossover was performed by plating these strains on LB agar without NaCl supplemented with 10% sucrose. Mutants were then confirmed by PCR analyses and whole-genome sequencing.

The *invL* complementation construct was generated by amplifying the putative promoter region (∼300 bp upstream) along with the *invL* open reading frame (primers: 5’ G636 *fdeC*KO Comp and 3’ G636 *fdeC*KO Comp-His6 v2) and the pUC18T-miniTn7T-Apr vector (primers: Tn7 linear Fwd-His6 and Tn7 liner Rev) (116). These amplicons were then assembled, generating pUC18T-miniTn7T-Apr::G636 *invL*KO comp. To generate the *gfp* integration construct, the *gfp* cassette was amplified from PB-FLuc+GFPd2 (primers: 5’ d2EGFP for pUC18T-mTn7 and 3’ d2EGFP for pUC18T-mTn7) and pUC18T-mTn7-Apr was amplified (primers: 5’ pUC18T-mTn7 for d2EGFP and 3’ pUC18T-mTn7 for d2EGFP). These fragments were then assembled, generating pUC18T-miniTn7T-Apr::*gfpd2*. pUC18T-miniTn7T-Apr::G636 *invL*KO comp and pUC18T-miniTn7T-Apr::*gfpd2* were introduced into G636 Δ*invL* and G636, respectively, using a four-parental conjugation technique, as previously described (116–119). Selection was achieved using LB supplemented with apramycin and chloramphenicol, and insertion of the respective fragments at the mTn7 site in the resulting G636 *invL*^+^ and G636-*gfp* strains was confirmed by PCR analyses.

### Antibiotic susceptibility assays

MIC analyses were performed using a two-fold broth dilution microtiter assay similar to previously described protocols (112, 120, 121). Briefly, overnight cultures were sub-cultured at 0.05 Abs600 and grown for 3 h shaking at 37 hpi. Mid-exponential growth phase cultures were then inoculated at 0.01 Abs600 into a 96-well microtiter plate (Corning Inc, Corning, NY) containing two-fold decreasing dilutions of the indicated antibiotics. Plates were then incubated at 37°C with shaking for 24 h. The MIC was defined as less than 10% of the Abs600 of an untreated control.

### Statistical methods

All statistical analyses were performed using GraphPad Prism version 9, and *P* values of <0.05 were considered statistically significant. When normally distributed, data sets were analyzed with Unpaired Student’s *t*-tests (comparing two samples), one-way <u>an</u>alysis <u>o</u>f <u>va</u>riance (ANOVA) with Tukey’s test for multiple comparisons (comparing more than two samples), or two-way ANOVA with Tukey’s test for multiple comparisons (comparing more than two samples with two independent variables). For non-normally distributed data sets, the Mann-Whitney *U* test (comparing two samples) or the Kruskal Wallis *H* test with Dunn’s test for multiple comparisons (comparing more than two samples) was used.

## Acknowledgements

This work was supported by funding to M.F.F. (R01AI166359 - CHECK), C.J.L. (T32AI007172), and C.B.L (R01AI137062) through the National Institute of Allergy and Infectious Diseases of the National Institutes of Health. JM was supported through The American Association of Immunologists Careers in Immunology Fellowship Program and The Pediatric Cardiovascular and Pulmonary Research Training Program (5T32HL125241-07). The modern respiratory isolates used in this study, G636 (strain 3689) and G654 (strain 6919), were collected by the CDC-funded Georgia <u>E</u>merging Infections <u>P</u>rogram’s (EIP) <u>Mu</u>lti-site <u>G</u>ram-Negative <u>S</u>urveillance Initiative (MuGSI) and kindly provided by Sarah Satola. We also acknowledge Jennifer Philips, Jacco Boon, and Gayan Bamunuarachchi for thoughtful discussion about the manuscript. We thank Wandy Beatty and the Washington University School of Medicine Molecular Microbiology Imaging Facility for microscopy assistance, Alma Johnson of the Washington University Center for Reproductive Health Sciences Histocore for lung tissue slide mounting and staining assistance, and De Chen of the Washington University Center for Cellular Imaging for assistance with the Zeiss AxioScan Z1. Finally, we thank Dakota Hall for technical assistance with experiments.

## Tables

**Table S1.** Cytokine analysis at 4 h, 2 d, and 7 d post-intranasal infection with G636.

| Cytokine<sup>a,b</sup> | WT |  |  | *tlr4* mutant |  |  |
| --- | --- | --- | --- | --- | --- | --- |
|  | 10<sup>5</sup> G636 | 10<sup>8</sup> G636 | Mock | 10<sup>5</sup> G636 | 10<sup>8</sup> G636 | Mock |
| **4 h** |  |  |  |  |  |  |
| IL-23 | 23.26 (10.14) | 38.53 (6.36) | 10.67 (7.09) | 22.57 (13.26) | 32.61 (3.61) | 15.90 (15.90) |
| IL-1α | 58.57 (18.27) | 323.11 (75.42)<sup>#, \$</sup> | 2.5 (0.43) | 2.77 (0.47) | 429.19 (90.22)<sup>#, \$</sup> | 1.49 (0.13) |
| IFN-γ | 0.77 (0.36) | 5.13 (1.30)<sup>#, \$</sup> | 0.00 (0.00) | 0.23 (0.23) | 4.34 (0.39)<sup>#, \$</sup> | 0.00 (0.00) |
| TNF-α | 2219.55 (214.72) | 14888.73 (311.27)<sup>#, \$</sup> | 108.38 (66.66) | 49.51 (6.47) | 12802.47 (3073.56)<sup>#, \$</sup> | 12.82 (3.58) |
| MCP-1 | 0.00 (0.00) | 65.73 (15.07)<sup>#, \$</sup> | 0.00 (0.00) | 0.00 (0.00) | 65.57 (8.23)<sup>#, \$</sup> | 0.00 (0.00) |
| IL-12p70 | 0.00 (0.00) | 1.43 (1.43) | 0.00 (0.00) | 0.00 (0.00) | 0.00 (0.00) | 0.00 (0.00) |
| IL-1β | 9.50 (1.95) | 46.26 (10.54)<sup>#, \$</sup> | 0.00 (0.00) | 0.00 (0.00) | 56.58 (12.58)<sup>#, \$</sup> | 0.00 (0.00) |
| IL-10 | 0.00 (0.00) | 4.18 (4.18) | 0.00 (0.00) | 0.00 (0.00) | 2.04 (2.04) | 0.00 (0.00) |
| IL-6 | 910.41 (172.71) | 7885.57 (1645.56)<sup>#, \$</sup> | 18.19 (5.67) | 8.09 (2.69) | 6334.72 (1218.67)<sup>#, \$</sup> | 2.70 (2.70) |
| IL-27 | 0.00 (0.00) | 85.90 (13.54)<sup>#, \$</sup> | 0.00 (0.00) | 0.00 (0.00) | 42.77 (24.83)<sup>\$</sup> | 0.00 (0.00) |
| IL-17A | 1.05 (0.74) | 8.36 (2.46)<sup>#, \$</sup> | 0.00 (0.00) | 0.00 (0.00) | 4.66 (0.39)<sup>#, \$</sup> | 0.00 (0.00) |
| IFN-β | 0.00 (0.00) | 29.75 (17.23)<sup>*, #, \$</sup> | 0.00 (0.00) | 0.00 (0.00) | 0.00 (0.00) | 0.00 (0.00) |
| GM-CSF | 96.65 (13.37)<sup>*, #</sup> | 55.42 (7.19)<sup>#, \$</sup> | 0.00 (0.00) | 0.00 (0.00) | 47.41 (6.96)<sup>#, \$</sup> | 0.00 (0.00) |
| **2 d** |  |  |  |  |  |  |
| IL-23 | 19.93 (9.61) | 27.05 (8.76) | 44.09 (21.87) | 26.04 (11.37) | 11.35 (5.56) | 72.50 (36.30) |
| IL-1α | 1.14 (0.19) | 157.28 (58.45)<sup>*, #, \$</sup> | 1.13 (0.20) | 2.00 (0.50) | 25.68 (6.15) | 19.64 (18.02) |
| IFN-γ | 2.20 (0.98) | 212.26 (54.96)<sup>*, #, \$</sup> | 0.00 (0.00) | 3.70 (2.48) | 4.96 (2.48) | 0.00 (0.00) |
| TNF-α | 1.85 (0.82) | 592.26 (154.11)<sup>*, #, \$</sup> | 1.03 (0.60) | 19.79 (6.79) | 52.20 (8.86) | 5.76 (4.31) |
| MCP-1 | 0.00 (0.00) | 160.30 (14.27)<sup>*, #, \$</sup> | 0.00 (0.00) | 0.00 (0.00) | 33.52 (6.37)<sup>#, \$</sup> | 0.00 (0.00) |
| IL-12p70 | 0.00 (0.00) | 46.17 (17.63)<sup>*, #, \$</sup> | 0.00 (0.00) | 0.00 (0.00) | 0.00 (0.00) | 0.00 (0.00) |
| IL-1β | 1.06 (1.06) | 12.35 (2.80)<sup>*, #, \$</sup> | 0.00 (0.00) | 0.00 (0.00) | 0.00 (0.00) | 0.00 (0.00) |
| IL-10 | 0.00 (0.00) | 0.00 (0.00) | 0.00 (0.00) | 0.00 (0.00) | 0.00 (0.00) | 0.00 (0.00) |
| IL-6 | 0.00 (0.00) | 1179.63 (323.29)<sup>*, #, \$</sup> | 0.00 (0.00) | 0.00 (0.00) | 20.45 (3.83) | 4.29 (4.29) |
| IL-27 | 0.00 (0.00) | 137.76 (81.43)<sup>*, #, \$</sup> | 0.00 (0.00) | 0.00 (0.00) | 0.00 (0.00) | 0.00 (0.00) |
| IL-17A | 0.42 (0.42) | 25.31 (10.22)<sup>*, #, \$</sup> | 0.00 (0.00) | 0.00 (0.00) | 0.00 (0.00) | 0.00 (0.00) |
| IFN-β | 0.00 (0.00) | 0.00 (0.00) | 0.00 (0.00) | 0.00 (0.00) | 0.00 (0.00) | 0.00 (0.00) |
| GM-CSF | 0.00 (0.00) | 0.00 (0.00) | 0.00 (0.00) | 0.00 (0.00) | 0.00 (0.00) | 0.00 (0.00) |
| **7 d** |  |  |  |  |  |  |
| IL-23 | 41.43 (14.43) | 30.14 (19.51) | 20.66 (20.66) | 9.07 (5.32) | 29.77 (12.87) | 12.69 (12.69) |
| IL-1α | 1.05 (0.18) | 41.50 (23.66) | 1.62 (0.92) | 5.54 (4.27) | 4.60 (2.50) | 1.79 (0.85) |
| IFN-γ | 0.00 (0.00) | 53.36 (40.45)<sup>\$</sup> | 0.00 (0.00) | 0.00 (0.00) | 0.00 (0.00) | 0.00 (0.00) |
| TNF-α | 0.00 (0.00) | 93.32 (36.14)<sup>*, #, \$</sup> | 0.00 (0.00) | 3.17 (3.17) | 2.41 (1.54) | 0.00 (0.00) |
| MCP-1 | 0.00 (0.00) | 14.83 (14.83) | 0.00 (0.00) | 0.00 (0.00) | 0.00 (0.00) | 0.00 (0.00) |
| IL-12p70 | 0.00 (0.00) | 1.54 (1.54) | 0.00 (0.00) | 0.00 (0.00) | 0.00 (0.00) | 0.00 (0.00) |
| IL-1β | 0.00 (0.00) | 1.43 (1.43) | 0.00 (0.00) | 0.00 (0.00) | 0.00 (0.00) | 0.00 (0.00) |
| IL-10 | 0.00 (0.00) | 6.84 (6.84) | 0.00 (0.00) | 0.00 (0.00) | 0.00 (0.00) | 0.00 (0.00) |
| IL-6 | 0.00 (0.00) | 232.41 (182.45) | 0.00 (0.00) | 5.19 (5.19) | 16.72 (16.72) | 0.00 (0.00) |
| IL-27 | 0.00 (0.00) | 26.87 (26.87) | 0.00 (0.00) | 0.00 (0.00) | 0.00 (0.00) | 0.00 (0.00) |
| IL-17A | 0.00 (0.00) | 13.67 (11.48) | 0.00 (0.00) | 0.00 (0.00) | 0.00 (0.00) | 0.00 (0.00) |
| IFN-β | 0.00 (0.00) | 0.00 (0.00) | 0.00 (0.00) | 0.00 (0.00) | 0.00 (0.00) | 0.00 (0.00) |
| GM-CSF | 0.00 (0.00) | 0.00 (0.00) | 0.00 (0.00) | 0.00 (0.00) | 0.00 (0.00) | 0.00 (0.00) |

<sup>a</sup>Mean pg/ml (SEM) from two independent experiments at each timepoint is displayed.
<sup>b</sup>\* $P < 0.05$ relative to *tlr4* mutant at same inoculum; # $P < 0.05$ relative to mock in same mouse strain. \$ $P < 0.05$ relative to 10<sup>5</sup> inoculum in same mouse strain. Two-way ANOVA, Tukey's test for multiple comparisons. Significant differences are also highlighted in green.

**Table S2.** MICs for *A. baumannii* strains G636 and G654.

| **Antibiotic** | **G636**<br>**(Resistant/Sensitive)** | **G654**<br>**(Resistant/Sensitive)** | **Clinical**<br>**Breakpoint<sup>a</sup>** |
| --- | --- | --- | --- |
| **Imipenem** | >256 µg/ml (Resistant) | >256 µg/ml (Resistant) | 2 µg/ml |
| **Ampicillin** | >256 µg/ml (N/A) | >256 µg/ml (N/A) | N/A<sup>c</sup> |
| **Ciprofloxacin** | >256 µg/ml (Resistant) | >256 µg/ml (Resistant) | 1 µg/ml |
| **Levofloxacin** | 32 µg/ml (Resistant) | 128 µg/ml (Resistant) | 2 µg/ml |
| **Colistin** | 1 µg/ml (Intermediate)<sup>b</sup> | 8 µg/ml (Resistant) | 2 µg/ml<sup>b</sup> |
| **Polymyxin B** | 2 µg/ml (Intermediate)<sup>b</sup> | 4 µg/ml (Resistant) | 2 µg/ml<sup>b</sup> |
| **Tigecycline** | 2 µg/ml (N/A)<sup>c</sup> | 1 µg/ml (N/A)<sup>c</sup> | N/A<sup>c</sup> |
| **Gentamicin** | >256 µg/ml (Resistant) | 2-4 µg/ml (Sensitive) | 4 µg/ml |
| **Apramycin** | 16 µg/ml (N/A)<sup>c</sup> | 16 µg/ml (N/A)<sup>c</sup> | N/A<sup>c</sup> |

<sup>a</sup>Clinical breakpoints are according to the Clinical and Laboratory Standard Institute (CLSI) M100 Performance Standards for Antimicrobial Susceptibility Testing 30<sup>th</sup> Edition (122).
<sup>b</sup> A “sensitive” breakpoint is not available for colistin or polymyxin B from the CLSI. Strains with MICs of less than or equal to 2 µg/ml are considered to have “intermediate resistance.”
<sup>c</sup>The clinical breakpoint has not been defined for ampicillin, tigecycline, and apramycin by the CLSI.

**Table S3.**
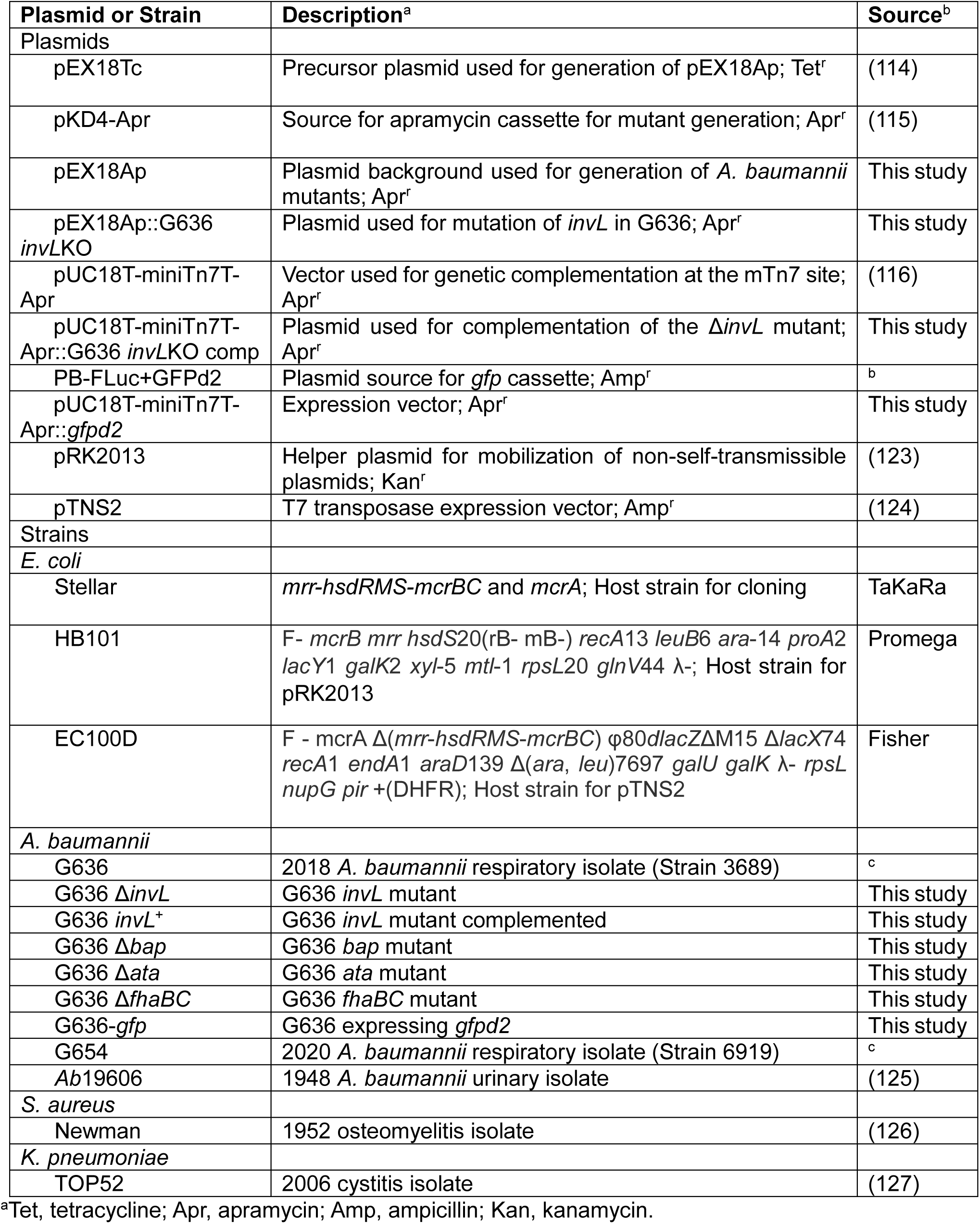
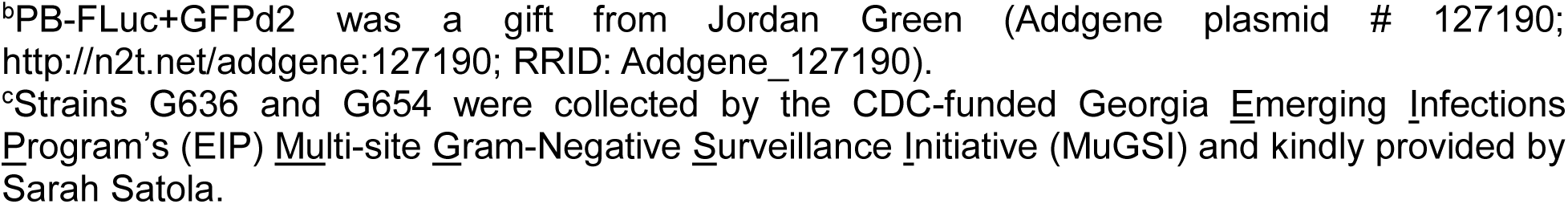
Plasmids and strains used in this study.

| Plasmid or Strain | Description<sup>a</sup> | Source<sup>b</sup> |
| --- | --- | --- |
| Plasmids |  |  |
| pEX18Tc | Precursor plasmid used for generation of pEX18Ap; Tet<sup>r</sup> | (114) |
| pKD4-Apr | Source for apramycin cassette for mutant generation; Apr<sup>r</sup> | (115) |
| pEX18Ap | Plasmid background used for generation of *A. baumannii* mutants; Apr<sup>r</sup> | This study |
| pEX18Ap::G636 *invLKO* | Plasmid used for mutation of *invL* in G636; Apr<sup>r</sup> | This study |
| pUC18T-miniTn7T-Apr | Vector used for genetic complementation at the mTn7 site; Apr<sup>r</sup> | (116) |
| pUC18T-miniTn7T-Apr::G636 *invLKO* comp | Plasmid used for complementation of the $\Delta invL$ mutant; Apr<sup>r</sup> | This study |
| PB-FLuc+GFPd2 | Plasmid source for *gfp* cassette; Amp<sup>r</sup> | <sup>b</sup> |
| pUC18T-miniTn7T-Apr::gfpd2 | Expression vector; Apr<sup>r</sup> | This study |
| pRK2013 | Helper plasmid for mobilization of non-self-transmissible plasmids; Kan<sup>r</sup> | (123) |
| pTNS2 | T7 transposase expression vector; Amp<sup>r</sup> | (124) |
| Strains |  |  |
| *E. coli* |  |  |
| Stellar | *mrr-hsdRMS-mcrBC* and *mcrA*; Host strain for cloning | TaKaRa |
| HB101 | F- *mcrB mrr hsdS20*(rB- mB-) *recA13 leuB6 ara-14 proA2 lacY1 galK2 xyl-5 mtl-1 rpsL20 glnV44* λ-; Host strain for pRK2013 | Promega |
| EC100D | F - *mcrA* $\Delta$(*mrr-hsdRMS-mcrBC*) ϕ80*dlacZ*ΔM15 Δ*lacX74 recA1 endA1 araD139* $\Delta$(*ara, leu*)7697 *galU galK* λ- *rpsL nupG pir* +(DHFR); Host strain for pTNS2 | Fisher |
| *A. baumannii* |  |  |
| G636 | 2018 *A. baumannii* respiratory isolate (Strain 3689) | <sup>c</sup> |
| G636 $\Delta invL$ | G636 *invL* mutant | This study |
| G636 *invL*<sup>+</sup> | G636 *invL* mutant complemented | This study |
| G636 $\Delta bap$ | G636 *bap* mutant | This study |
| G636 $\Delta ata$ | G636 *ata* mutant | This study |
| G636 $\Delta fhaBC$ | G636 *fhaBC* mutant | This study |
| G636-*gfp* | G636 expressing *gfpd2* | This study |
| G654 | 2020 *A. baumannii* respiratory isolate (Strain 6919) | <sup>c</sup> |
| Ab19606 | 1948 *A. baumannii* urinary isolate | (125) |
| *S. aureus* |  |  |
| Newman | 1952 osteomyelitis isolate | (126) |
| *K. pneumoniae* |  |  |
| TOP52 | 2006 cystitis isolate | (127) |

<sup>a</sup>Tet, tetracycline; Apr, apramycin; Amp, ampicillin; Kan, kanamycin.
<sup>b</sup>PB-FLuc+GFPd2 was a gift from Jordan Green (Addgene plasmid # 127190; http://n2t.net/addgene:127190; RRID: Addgene\_127190).
<sup>c</sup>Strains G636 and G654 were collected by the CDC-funded Georgia Emerging Infections Program's (EIP) Multi-site Gram-Negative Surveillance Initiative (MuGSI) and kindly provided by Sarah Satola.

**Table S4.**
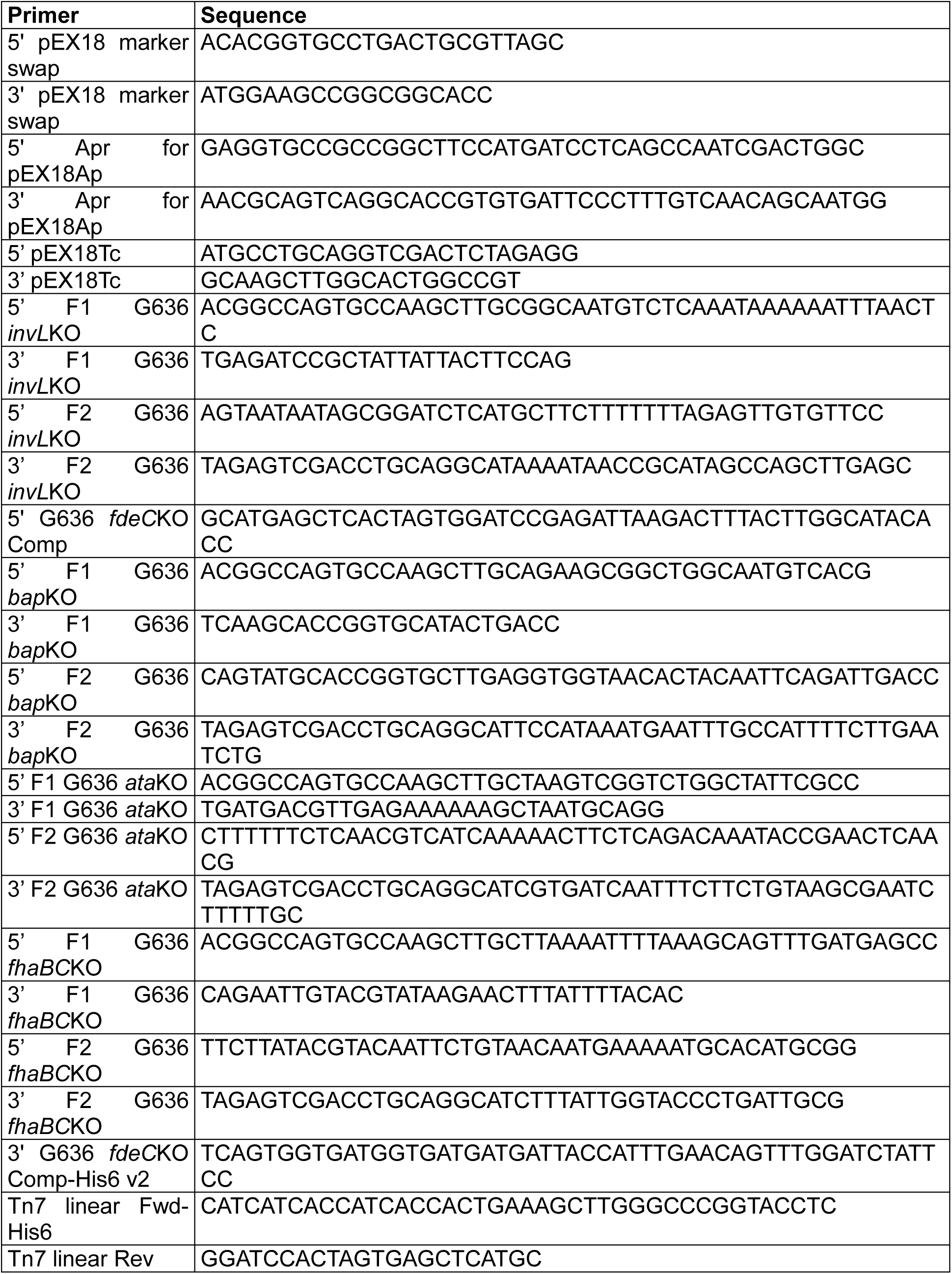
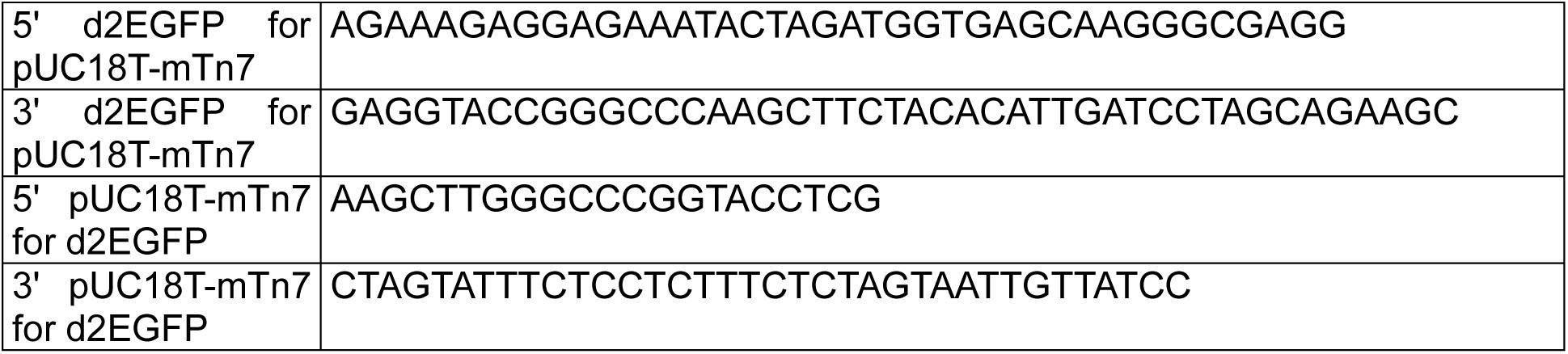
Primers used in this study.

## Figure Legends

**Figure S1.**
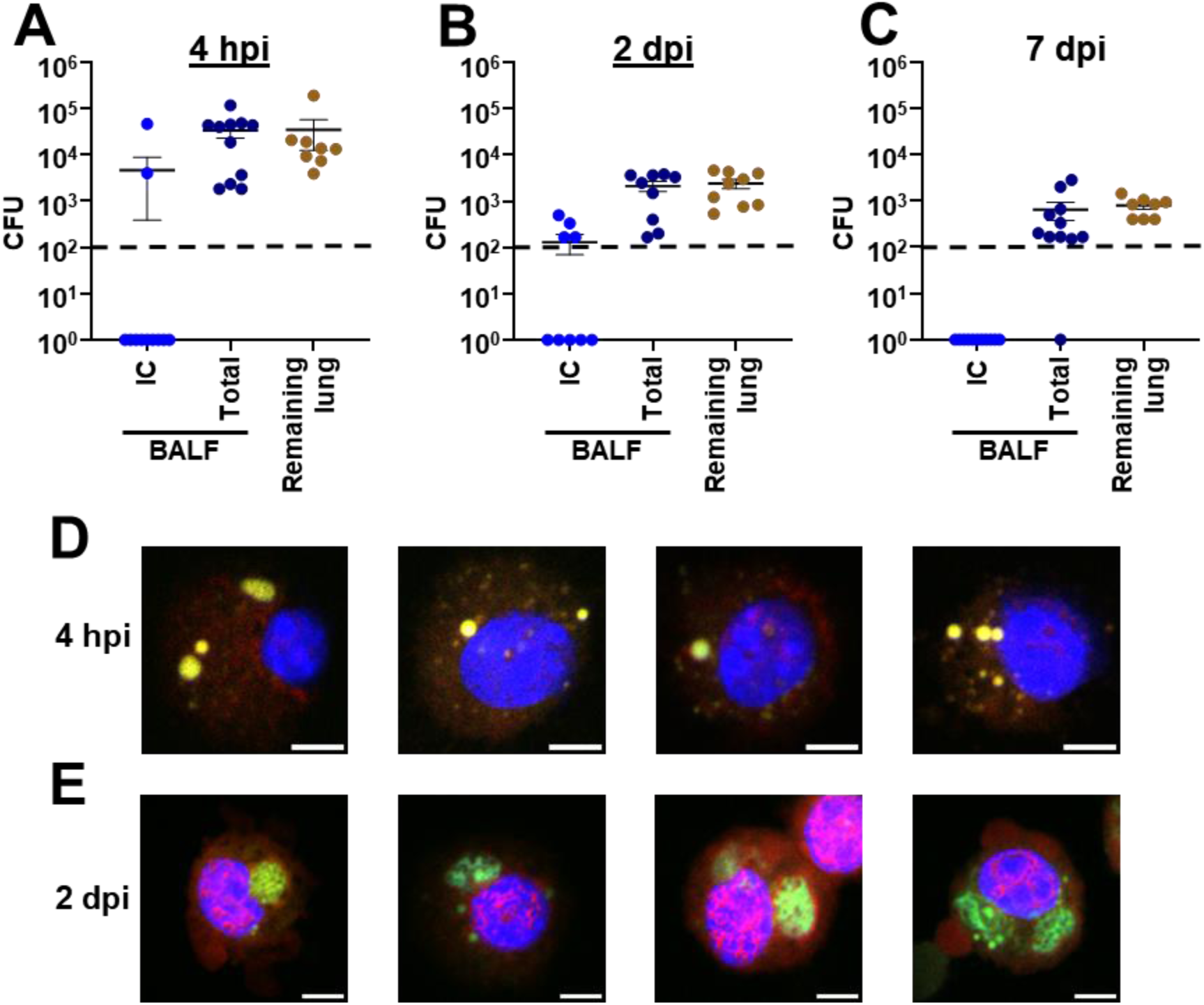
Intracellular *A. baumannii* are detectable in BALF at early timepoints in the chronic respiratory infection model. Groups of C3H/HeJ (*tlr4* mutant) mice were intranasally inoculated with 10^5^ G636, and BALF was collected at 4 hpi (A), 2 dpi (B), and 7 dpi (C) and either treated with 50 µg/ml colistin or mock-treated. Following, bacterial CFU in the treated (intra<u>c</u>ellular; IC) and mock treated (total) BALF, as well as in the remaining lungs following BALF collection, were enumerated by serial dilution plating. The horizontal line represents the mean, and the SEM is indicated by error bars. Shown are the results from at least two independent experiments. C3H/HeJ (*tlr4* mutant) mice were infected with G636 expressing *gfp*, and, at these same timepoints, BALF was collected, and host cells were isolated and stained with DAPI (blue) and phalloidin (red). Intracellular bacteria were identified by microscopy at 4 hpi (D) and 2 dpi (E). Shown are representative images from independent samples from at least two biological replicates. Scale bar = 5 µm.

**Figure S2.**
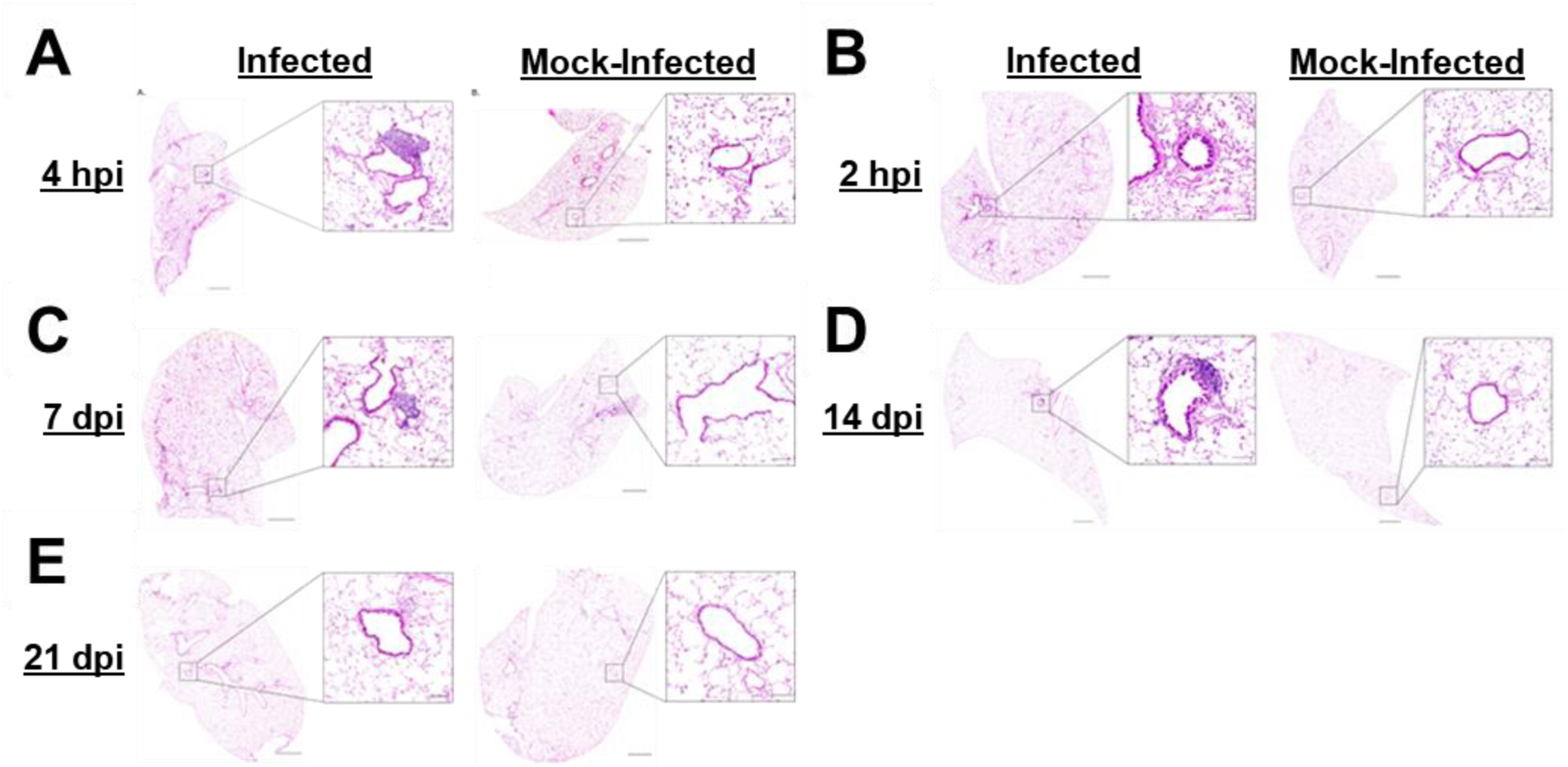
The chronic respiratory infection model results in lung pathology during infection. Groups of C3H/HeJ (*tlr4* mutant) mice were inoculated with 10^5^ G636 or mock-inoculated with PBS, and at 4 hpi (A), 2 dpi (B), 7 dpi (C), 14 dpi (D), and 21 dpi (E), lungs slices were prepared and H&E stained. Shown are representative images from each timepoint. Lung slice scale bar: 1000 µm; Inset scale bar: 100 µm.

**Figure S3.**
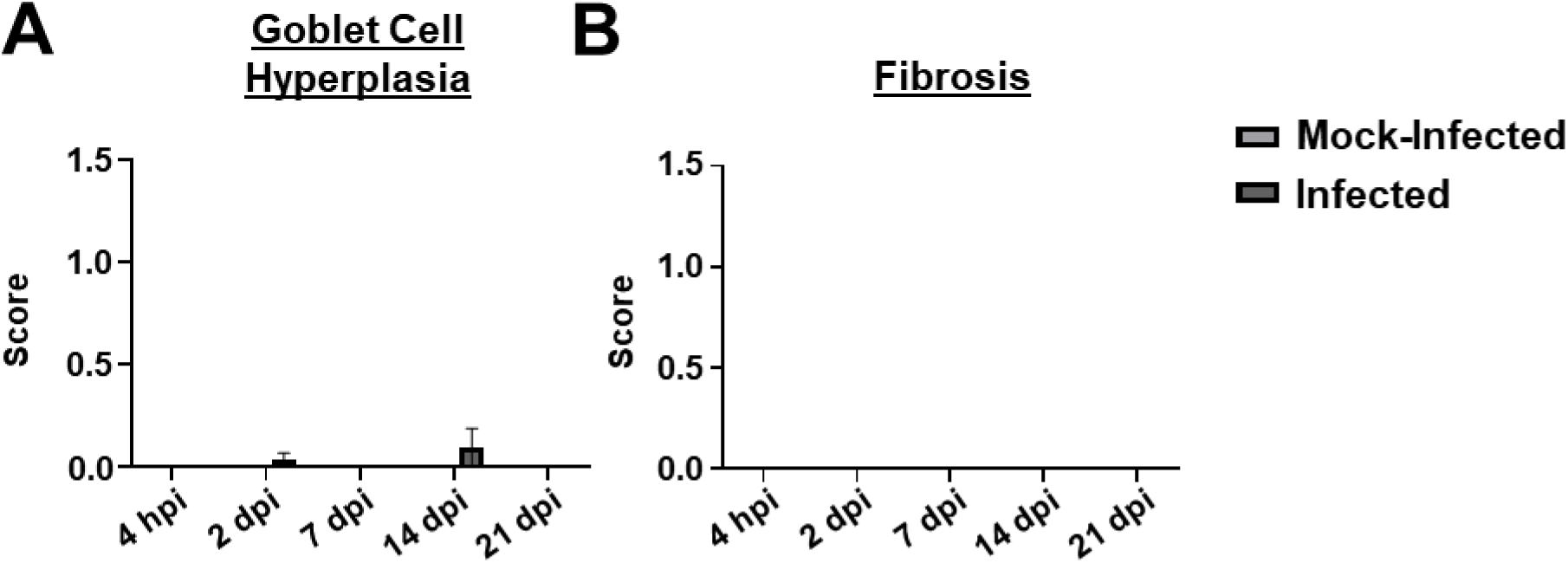
The chronic respiratory infection model does not cause goblet cell hyperplasia or fibrosis. Groups of C3H/HeJ (*tlr4* mutant) mice were inoculated with 10^5^ G636 or mock-inoculated with PBS, and at 4 hpi, 2 dpi, 7 dpi, 14 dpi, and 21 dpi, lungs slices were prepared, H&E stained, and scored for goblet cell hyperplasia (A) and fibrosis (B). The mean is shown on the graph, and the SEM is indicated by error bars. Unpaired Student’s *t*-test.

**Figure S4.**
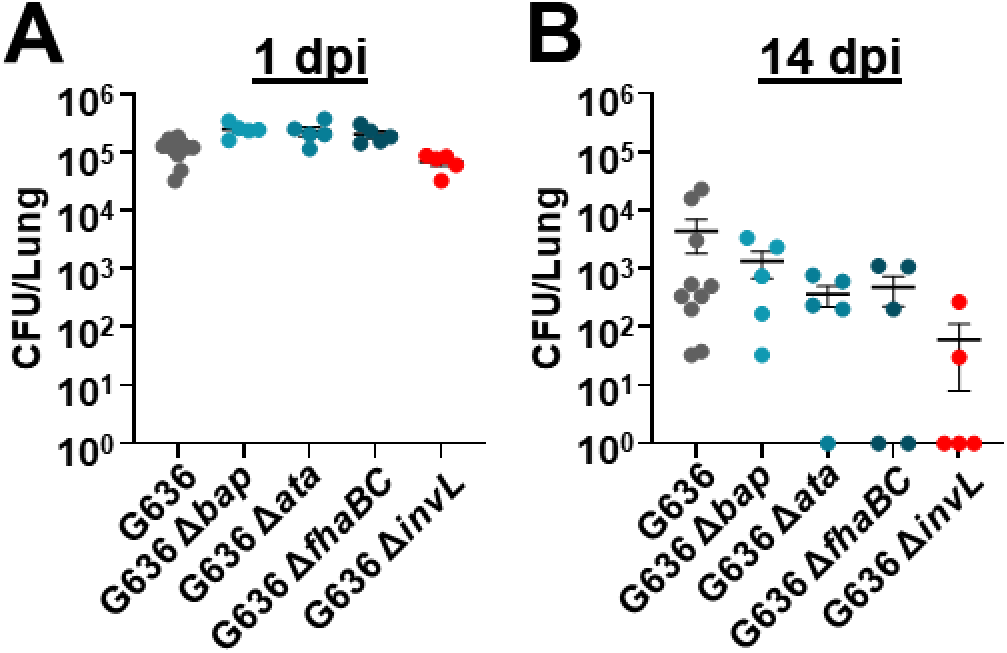
Testing of G636 adhesin mutants in the chronic respiratory infection model reveals a potential role for InvL in bacterial persistence. Groups of C3H/HeJ (*tlr4* mutant) mice were intranasally inoculated with 10^5^ G636, G636 Δ*bap*, G636 Δ*ata*, G636 Δ*fhaBC*, and G636 Δ*invL*. 1 (A) and 14 (B) dpi, mice were sacrificed, and CFU in the lungs were quantified. Each data point represents an individual mouse, the horizontal line represents the mean, and the SEM is indicated by error bars. Shown are results from single experiments for each strain.

**Figure S5.**
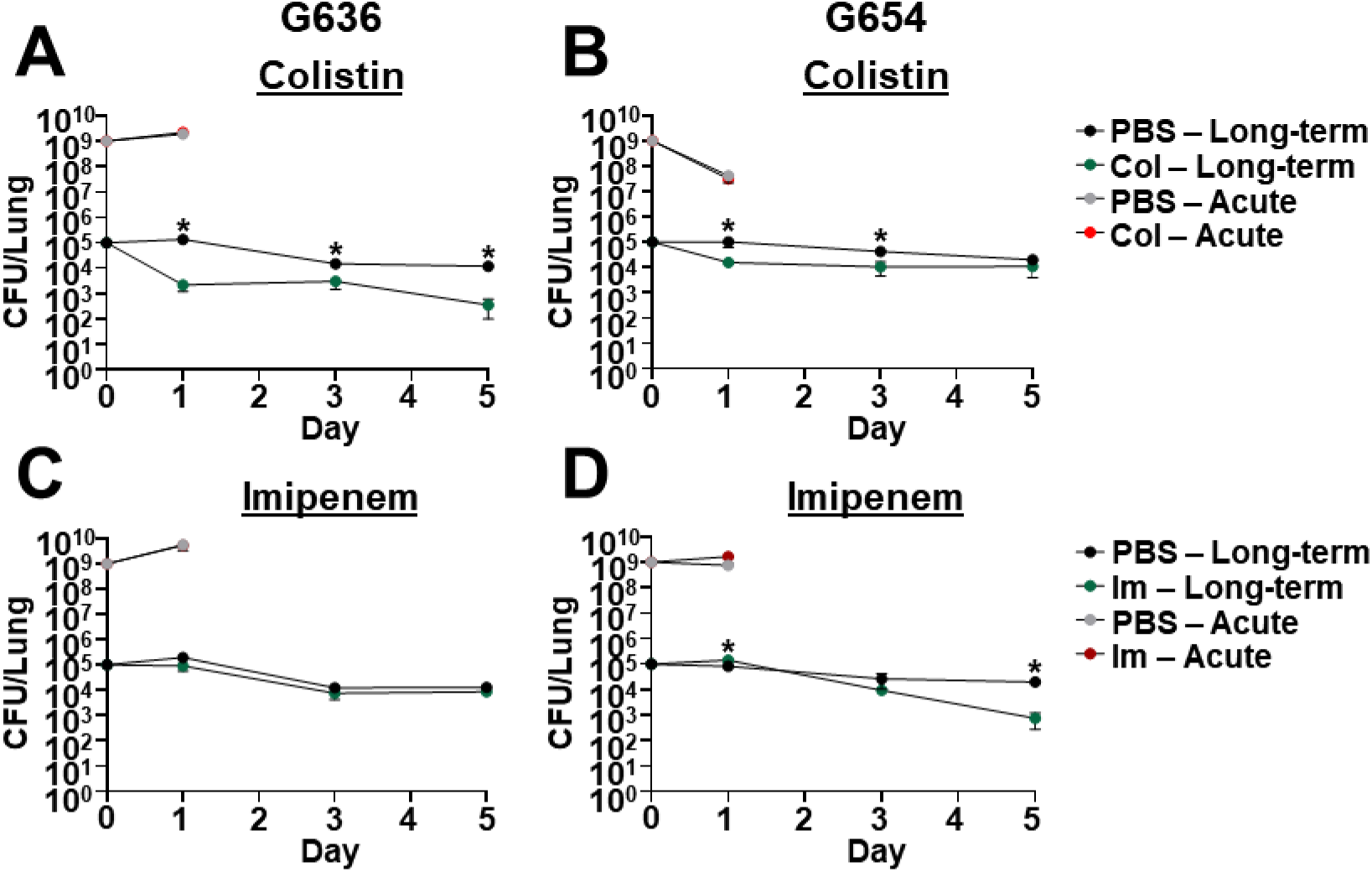
The chronic respiratory infection model can be used to study outcomes of antibiotic treatment. Groups of C3H/HeJ (*tlr4* mutant) mice were infected with 10^5^ G636 (A, C) or 10^5^ G654 (B, D) and sacrificed at 1, 3, and 5 dpi (long-term). Additionally, groups of C57Bl/6 mice were infected with 10^9^ G636 (A, C) or 10^9^ G654 (B, D) and sacrificed at 24 hpi (acute). Mice in both infection models were treated intraperitoneally treated with PBS or 5 mg/kg <u>col</u>istin (col) every 8 h (A, B) or PBS or 100 mg/kg <u>im</u>ipenem (im) every 12 h (C, D) with all treatments beginning 4 hpi. At each timepoint, CFU were quantified in the lungs. Shown are the results from at least two independent experiments, each data point represents an individual mouse, the horizontal line represents the mean, and the SEM is represented by error bars. \**P* < 0.05; Mann-Whitney *U* test.

**Figure S6.**
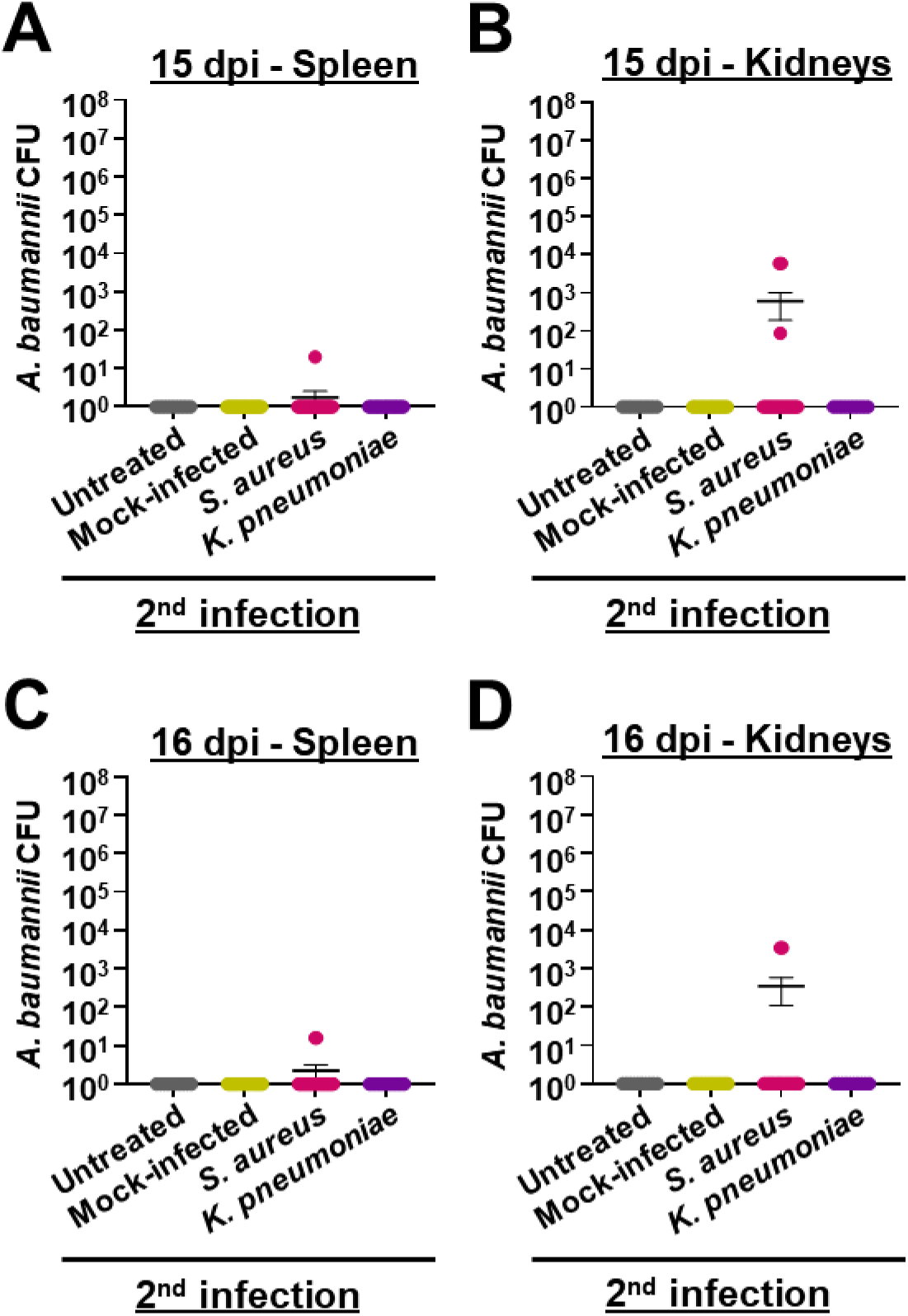
*S. aureus* secondary infection sometimes causes *A. baumannii* dissemination to the spleen and kidneys in the chronic respiratory infection model. C3H/HeJ (*tlr4* mutant) mice were intranasally inoculated with 10^5^ G636. At 14 days post-*A. baumannii* infection, groups of mice were either not inoculated (untreated), inoculated with PBS (mock-infected), infected with *S. aureus*, or infected with *K. pneumoniae*. Subsequently, on days 15 (A and B) and 16 (C and D) post-*A. baumannii* infection (1 and 2 days post-secondary infection), groups of mice were sacrificed, and *A. baumannii* CFU were quantified in the spleen (A and C), and kidneys (C and D). Each data point represents an individual mouse, the horizontal line represents the mean, and the SEM is indicated by error bars. Shown are results from at least 2 independent experiments. Significant differences were not detected; Kruskal-Wallis *H* test with Dunn’s test for multiple comparisons.

**Figure S7.**
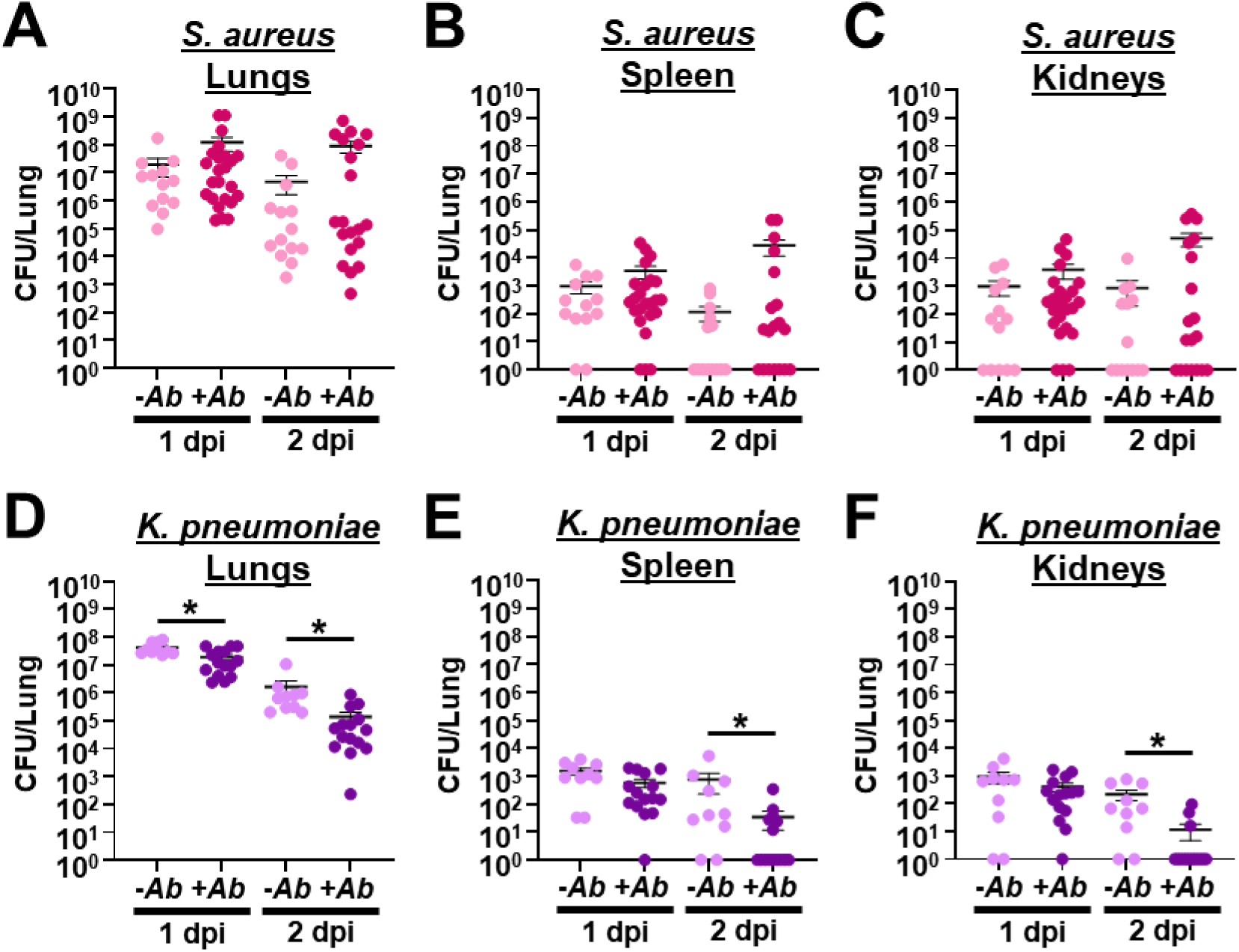
Ongoing *A. baumannii* pneumonia alters bacterial numbers following secondary infection with *S. aureus* and *K. pneumoniae*. C3H/HeJ (*tlr4* mutant) mice were either intranasally inoculated with 10^5^ G636 14 days prior to infection with *S. aureus* or *K. pneumoniae* (+*Ab*) or not infected prior to *S. aureus* or *K. pneumoniae* infection (-*Ab*). At 14 days post-*A. baumannii* infection, groups of mice were infected with *S. aureus* or *K. pneumoniae*. 1 and 2 dpi with *S. aureus* (A, B, and C) or *K. pneumoniae* (D, E, and F), mice were sacrificed, and these bacteria were quantified in the lung (A and D), spleen (B and E), and kidneys (C and F). Each data point represents an individual mouse, the horizontal line represents the mean, and the SEM is indicated by error bars. Shown are results from at least 2 independent experiments. \**P* < 0.05; Mann-Whitney *U* test.

